# Coupled Effects of Airway Deformation and Carrier Gas on Pulmonary Gas-Particle Flow Dynamics Using an Euler-Lagrange Method with Dynamic Mesh

**DOI:** 10.64898/2026.10.15.759419

**Authors:** Hang Yi, Yu Feng

**Author notes:** Corresponding Author: Address: 420 Engineering North, Stillwater, OK 74078, USA.

## Abstract

Physiologically realistic prediction of inhaled aerosol deposition in the airways requires simultaneous consideration of airway deformation kinematics and carrier-gas properties. This study developed a 3D deformable lung airway model (DLAM) using an experimentally validated computational fluid particle dynamics (CFPD) model coupled with a dynamic mesh method. Both the DLAM and a conventional static lung airway model (SLAM) were used to predict and compare inhaled gas-particle transport and deposition in a subject-specific mouth-to-generation 9 (G9) human respiratory system. To assess the significance of airway deformation, SLAM and DLAM were compared under identical 12 breaths per minute (BPM) inhalation-exhalation waveforms for both conventional air and heliox (70% He-30% O_2_), considering particle sizes of 1, 5, and 10 µm. The effects of carrier gas and breathing frequency were further evaluated using the more physiologically realistic DLAM, comparing air and heliox at 12, 16, and 20 BPM for the same particle sizes. Results showed that the influence of airway deformation on aerosol transport is strongly dependent on particle size. Differences between SLAM and DLAM were negligible for 1 µm particles. In contrast, for 5 and 10 µm particles, SLAM consistently underpredicted beyond-G9 delivery and overpredicted G0-G9 local deposition for both air and heliox. For example, for 10 µm particles at 12 BPM, accounting for airway deformation kinematics increased the beyond-G9 delivery fraction by an absolute 3.76% with air and 5.55% with heliox. After establishing the size-dependent deformation effect, DLAM was further used to evaluate the influence of carrier gas and breathing frequency on inhaled aerosol transport and deposition. Compared with air, heliox reduced turbulence kinetic energy, wall shear stress, and pressure drop, and enhanced beyond-G9 delivery of larger particles. Specifically, for 10 µm particles, the absolute increases in beyond-G9 delivery were 16.82%, 20.25%, and 22.90% at 12, 16, and 20 BPM, respectively. Increasing breathing frequency reduced distal delivery, particularly for 5 and 10 µm particles, whereas 1 µm particles maintained approximately 99% beyond-G9 penetration across all investigated conditions. These findings indicate that DLAM is essential for accurate regional dosimetry of higher-inertia aerosols, for which the static airway simplification can significantly bias predicted deposition and distal delivery.

## 1. Introduction

Inhalation therapy is widely used to deliver aerosolized medications via the pulmonary route, enabling high local concentration in the lung while minimizing systemic exposure (Feng et al., 2016; Feng et al., 2021; Kleinstreuer & Feng, 2013; Longest et al., 2019). However, because of complex airway morphology, current inhalation therapies can be inefficient at delivering drugs to target lung sites, e.g., small airways for chronic obstructive pulmonary disease (COPD) or asthma treatment (Labiris & Dolovich, 2003; Usmani et al., 2021). Indeed, off-target deposition can reduce the fraction of the administered dose reaching the intended region and may contribute to local or systemic adverse effects, thereby reducing the therapeutic efficiency of inhaled medications.

Achieving drug delivery optimization to the small airways requires a comprehensive understanding of inhaled aerosol transport and deposition dynamics, particularly the spatiotemporal distributions of carrier-gas flow and dispersed particles in the respiratory system. Obtaining such detailed *in vivo* information remains challenging, limited by imaging resolution and ethical reasons. As an alternative, experimentally validated computational fluid particle dynamics (CFPD) is a first principle based noninvasive and cost-effective methodology to unveil the underlying gas-particle flow dynamics (Feng et al., 2021; Longest et al., 2019). However, most current respiratory CFPD models assume static airway walls, even though the glottis and intrathoracic airways undergo continuous deformation during breathing. Such deformation kinematics alter the transient carrier gas flow field and gas-particle interactions, which may influence the accuracy of inhaled particle deposition predictions. Therefore, incorporating physiologically realistic airway deformation kinematics in the CFPD airway model (i.e., CFPD-based moving lung model) is essential for improving CFPD-based predictions of medicinal aerosol dosimetry and ensuring the accuracy of the CFPD results so that they can be employed to guide innovation in inhalation therapy for enhanced targeted drug delivery to designated lung sites.

One emerging strategy to optimize aerosolized medication delivery is selecting the carrier gas. The density and viscosity of the carrier gas influence the local airflow regime (i.e., lower turbulence intensity, less flow resistance, etc.), thereby affecting particle transport and deposition sites (Farooq et al., 2025; Farooq et al., 2023; Huang et al., 2026; Yi & Feng, 2026). Recent CFPD research also shows that replacing air with a low-density carrier gas can reduce turbulence-related particle losses and promote drug delivery to distal airways (Yi & Feng, 2026). Helium-oxygen (He-O_2_) mixtures, known as heliox, are of particular interest due to their substantially lower density than air, which can reduce gas flow resistance and turbulence in constricted or highly curved airways (Gemci et al., 2003; Hess et al., 2006). As a result, heliox has been investigated as a carrier gas for inhaled medications, which can potentially increase drug delivery to the peripheral lung. Corcoran and Gamard (2004) reviewed the aerosol and flow physics of heliox and discussed their potential to enhance peripheral aerosol delivery. Gemci et al. (2003) numerically reported reduced extrathoracic aerosol deposition with 70% He-30% O_2_ heliox compared with air. Subsequent studies show that aerosol delivery with heliox depends on the drug delivery system configuration, gas composition, and inhalation-exhalation conditions (Corcoran, 2021; Li et al., 2024). Farooq et al. (2023) employed a CFPD model to examine aerosol transport in air, 78% He-22% O_2_ heliox, and a heliox-air mixture in an idealized generation 3 (G3)-G5 tracheobronchial (TB) geometry. The study showed that heliox reduced turbulence and particle deposition in the G3-G5 geometry, which may enhance aerosol delivery to airways downstream to G5. Subsequently, Farooq et al. (2025) expanded the CFPD analysis to a realistic TB tree model extending from the trachea to G5, and similarly observed that heliox decreased pressure loss, turbulence, and regional particle deposition compared with air. Collectively, these studies establish carrier-gas selection as a potentially effective strategy to increase the drug delivery efficiency to small airways.

However, existing CFPD simulations for gas carrier effects have relied heavily on static airway models, which do not consider physiologically realistic airway deformation kinematics (Zhao et al., 2020; Zhao et al., 2021). In reality, airways are dynamic during the inhalation-exhalation process: (1) the glottis opens and closes, and (2) intrathoracic airways (i.e., the TB tree) expand and contract throughout the breathing cycle (Zhao et al., 2020; Zhao et al., 2021). Such deformation kinematics can alter the transient carrier-gas flow field and gas-particle interactions, which may affect the accuracy of inhaled particle deposition predictions. Therefore, incorporating physiologically realistic airway deformation kinematics in the CFPD airway model (i.e., CFPD-based moving lung model) is essential for improving CFPD-based predictions of medicinal aerosol dosimetry to ensure the accuracy of the CFPD results so that they can be employed for guiding innovation in inhalation therapy for enhanced targeted drug delivery to designated lung sites, via multiple carrier gases. Previous studies have developed regional deformation kinematics models and integrated them into CFPD models (Aghasafari & Pidaparti, 2018; Faizal et al., 2020; Gunatilaka et al., 2024; Jahed et al., 2025; Jing et al., 2026; Kolanjiyil & Kleinstreuer, 2017; Li et al., 2023; Malvè et al., 2011; Samanta et al., 2026; Seyfi et al., 2016; Subramaniam et al., 2017; Talaat & Xi, 2017; Wall & Rabczuk, 2008; Wall et al., 2010; Werner et al., 2008; Xi et al., 2018; Xia et al., 2010; Zhao et al., 2020; Zhao et al., 2021). These studies demonstrate that physiologically realistic deformation kinematics of airways can alter air-particle flow dynamics in human respiratory systems, although the reported effects are not uniform. Some investigations have identified substantial differences between rigid and deforming airways, while others, particularly at the alveolar scale, have reported relatively weak sensitivity of particle deposition to specific deformation modes.

Thus, a critical question remains unresolved. Widely used static CFPD airway models cannot be assumed to consistently overpredict or underpredict aerosol deposition across all conditions. The magnitude and direction of these differences may depend on subject-specific airway geometry features, inhaled particle sizes, breathing conditions, and carrier-gas properties. Such an uncertainty is essential when CFPD is used not only to characterize aerosol transport, but also to select or optimize a carrier gas for enhanced drug delivery to small and peripheral airways. To date, carrier-gas effects such as those associated with air and heliox have not been systematically evaluated using a physiologically deforming, subject-specific mouth-to-G9 airway model. Conversely, previous moving-airway studies have not established whether carrier-gas-mediated aerosol-targeting trends obtained with rigid-airway models remain valid when physiologic airway deformation is incorporated. Therefore, the coupled effects of airway deformation kinematics and carrier-gas properties on regional medicinal aerosol delivery to small airways remain insufficiently understood.

The present study **innovatively** addresses these two interconnected gaps and represents the first CFPD effort to systematically evaluate carrier-gas effects in a physiologically deforming, subject-specific mouth-to-G9 airway model. Specifically, a subject-specific 3D deformable lung airway model (DLAM) (see **Fig. 1**) was developed from the established in-house dynamic mesh method (Zhao et al., 2020; Zhao et al., 2021) and coupled with an Euler-Lagrange model (Feng et al., 2021; Feng et al., 2018). Initially, a conventional static lung airway model (SLAM) and the DLAM are directly compared under identical 12 breaths per minute (BPM) breathing conditions using both conventional air and 70% He-30% O_2_ heliox for particles with multiple aerodynamic diameters (i.e., *d_p_* = 1, 5, and 10 *μm*). This comparison assesses how neglecting airway deformation influences predicted regional aerosol dosimetry and whether this effect is consistent across particle sizes and carrier gases. Subsequently, the DLAM is used to systematically investigate carrier-gas effects at 12, 16, and 20 BPM for the same three particle sizes. Through this approach, the study evaluates both the significance of physiologically realistic airway deformation for CFPD-based aerosol dosimetry and the potential of low-density carrier gases to enhance medicinal aerosol delivery to the distal lung. High-resolution CFPD analysis includes spatiotemporal distribution of velocity magnitude, turbulence kinetic energy (TKE), wall shear stress (WSS), pressure distributions, particle residence time (PRT), localized deposition patterns, regional deposition fraction (RDF), and beyond-G9 delivery fraction.

**Fig. 1.**
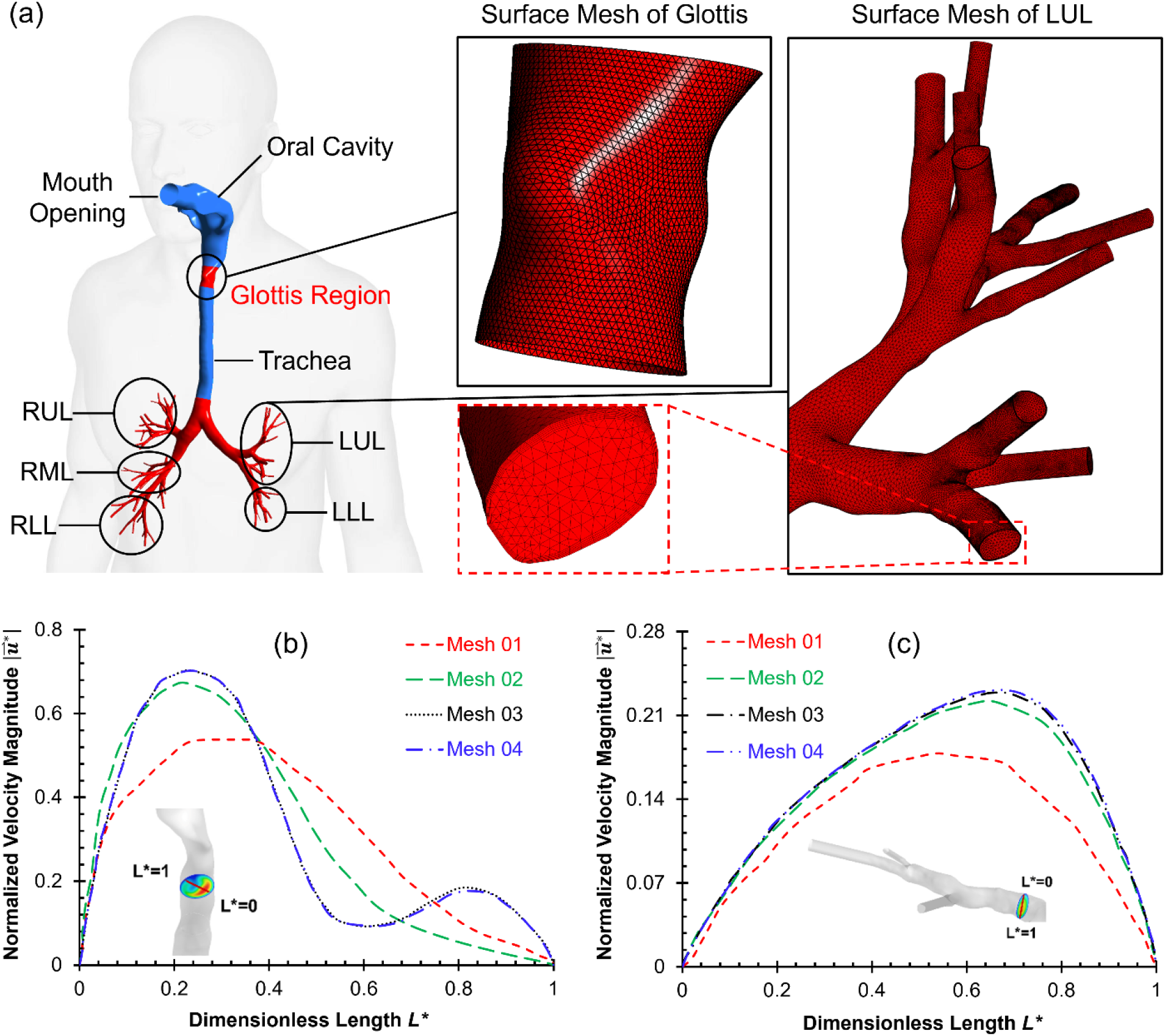
Geometry and mesh of the subject-specific mouth-to-G9 human respiratory system: (a) Geometry and tetrahedron-based mesh details with Mesh 03 selected as the final mesh, (b) Comparison of local normalized velocity magnitude along the selected line in trachea for mesh independence test, and (c) Comparison of local normalized velocity magnitude along the selected line G9 for mesh independence test.

The remainder of this paper is structured as follows. **Section 2** details the subject-specific airway geometry, mesh generation, governing equations, boundary conditions including prescribed airway deformation kinematics, and model validation. **Section 3** presents the results and discussion, beginning with a comparison of static and deforming elastic airway models (i.e., SLAM vs. DLAM), followed by an analysis of air vs. heliox, breathing pattern influence, and particle size effects using DLAM, concluding with a discussion of the potential physiological implications of heliox as gas carrier for inhalation therapy. **Section 4** summarizes the major conclusions, and **Section 5** discusses the limitations of the present study and future work.

## 2. Methodology

### 2.1 Geometry and mesh

**Figure 1 (a)** shows the 3D subject-specific human respiratory system employed in this study, extending from the mouth to up-to-G9 across five lobes. The five lobes are the left upper lobe (LUL), left lower lobe (LLL), right upper lobe (RUL), right middle lobe (RML), and right lower lobe (RLL), respectively. During reconstruction of the airway geometry, a widely used upper airway model (Worth Longest & Xi, 2008) was connected with a subject-specific TB tree reconstructed from CT scan data of a 30-year-old healthy male volunteer with no history of respiratory diseases. A 64-slice General Electric (GE) Healthcare computed tomography (CT) scanner was used to take images of 500 mm by 500 mm (i.e., 512×512 pixels on the plane) cross-sections with 2.5 mm image slice thickness. The total lung lobe volume, i.e., functional residual volume (FRC_WL_) is equal to 2714.98 ml, representing a body weight of 90.47 kg (30 ml/kg is the standard). The upper airway functional residual volume FRC_UA_ is 25.50 ml.

Tetrahedral meshing strategy was employed to generate CFPD meshes using Ansys Fluent Meshing 2025R2 (Synopsys Ansys Inc., Canonsburg, PA). The airflow domain was discretized using tetrahedral elements with 5 near-wall prism layers (see **Fig. 1 (a)**). Mesh independence tests were performed by comparing velocity distributions (i.e., normalized velocity magnitude 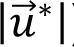) at two selected lines from mouth to generation G9 with four meshes (see **Figs. 1 (b) & (c)**). Specifically, the four successively refined meshes (i.e., Meshes 01-04) were generated, with Mesh 01 being the coarsest and Mesh 04 the finest. The final mesh (i.e., Mesh 03) was determined by refining the mesh until grid independence of the flow field solutions was achieved, that is, the relative 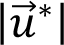 difference between Mesh 03 and finest mesh (Mesh 04) is smaller than 1%. Therefore, Mesh 03 is selected as the final mesh, which contains 2,796,297 nodes and 9,291,640 cells including 6,081,210 tetrahedrons and 3,210,430 prismatic cells.

### 2.2 Governing equations

In this study, two CFPD-based models were developed and employed: (1) a conventional static lung airway model (SLAM) using an experimentally validated Euler-Lagrange model (Feng et al., 2021; Feng et al., 2018), in which the airway walls were assumed to be stationary, and (2) a deformable lung airway model (DLAM), which incorporated the same Euler-Lagrange model (Feng et al., 2021; Feng et al., 2018) with a dynamic mesh method to simulate the physiologically realistic prescribed anisotropic airway deformation kinematics (i.e., glottis and TB tree motions shown in **Fig. 2**) (Zhao et al., 2020; Zhao et al., 2021).

**Fig. 2.**
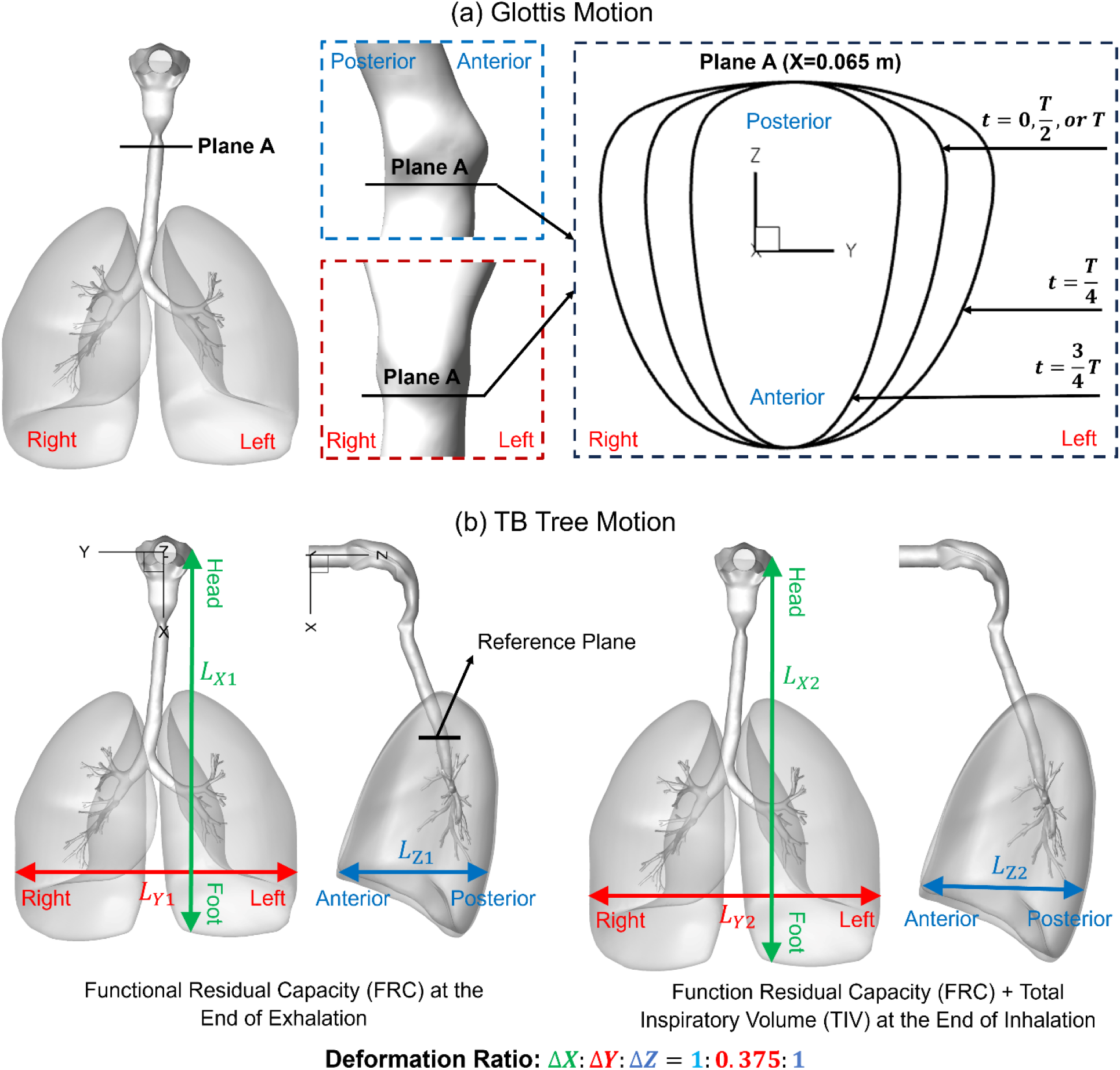
Airway deformation kinetics in the selected subject-specific human respiratory system: (a) Glottis motion, and (b) Anisotropic TB tree motion.

#### 2.2.1 Continuous phase (gas carriers)

For the continuous phase flow (i.e., air and heliox), an experimentally validated *k* -*ω* Transition Shear Stress Transport (SST) model (Chen et al., 2017; Liu et al., 2024; Zhang & Kleinstreuer, 2011) was employed to predict transient laminar-to-turbulence transitional flow characteristics through the airway over three complete breathing cycles. The breathing conditions were prescribed with an inhalation-to-exhalation duration ratio of 1:1 and a baseline tidal volume of 630 mL (see **Fig. 3**). The SLAM and DLAM were directly compared at 12 BPM, while the DLAM was further evaluated at 12, 16, and 20 BPM, respectively. The physical properties of air and heliox gas carriers are listed in **Table 1**, which are assumed to be incompressible and isothermal. Accordingly, the governing equations for the continuous gas phase are given as

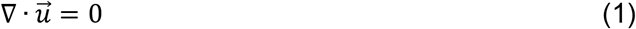

**Fig. 3.**
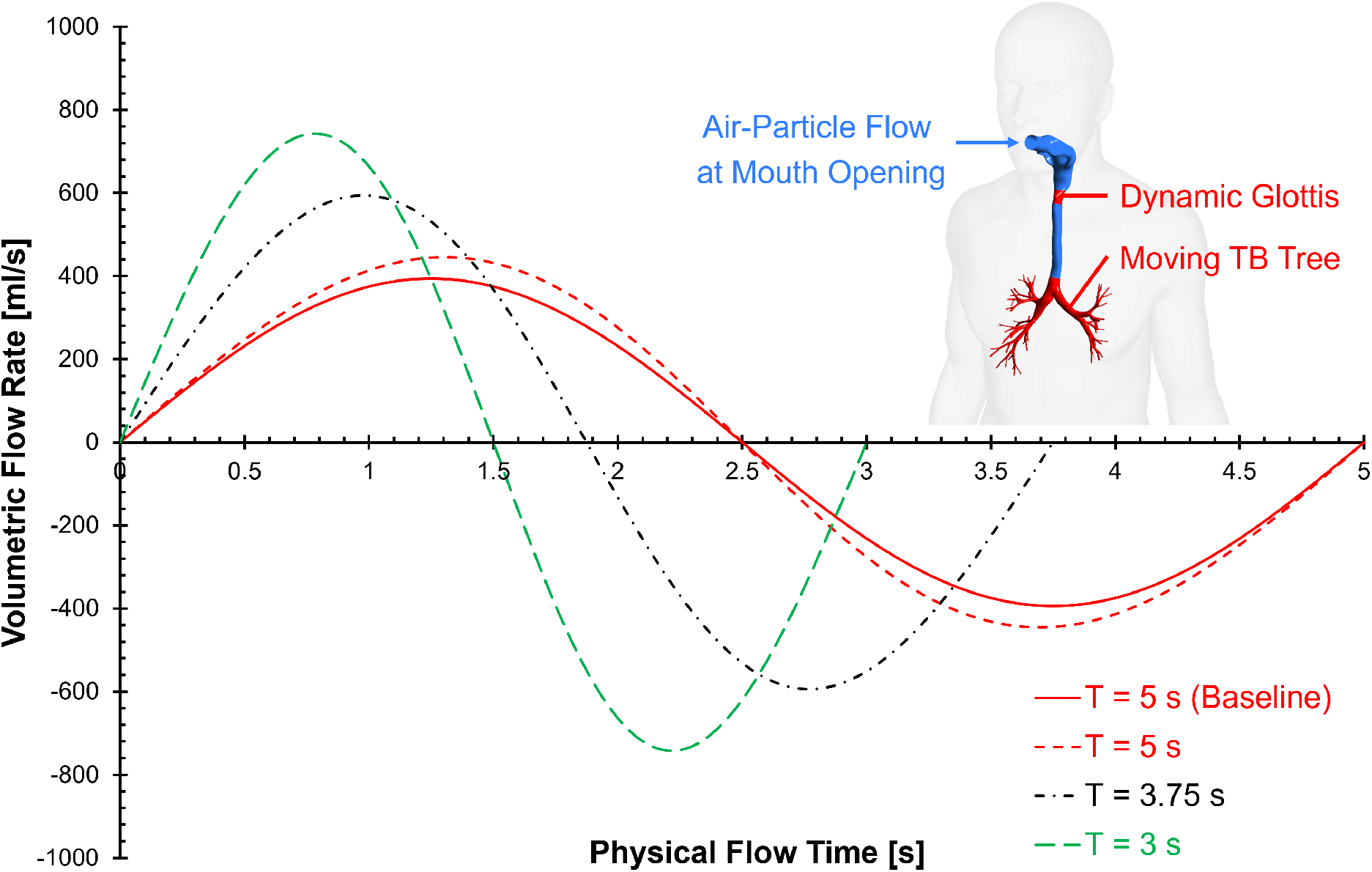
Breathing waveforms used in this study at different BPMs with a fixed tidal volume of 710 ml.

**Table 1:**
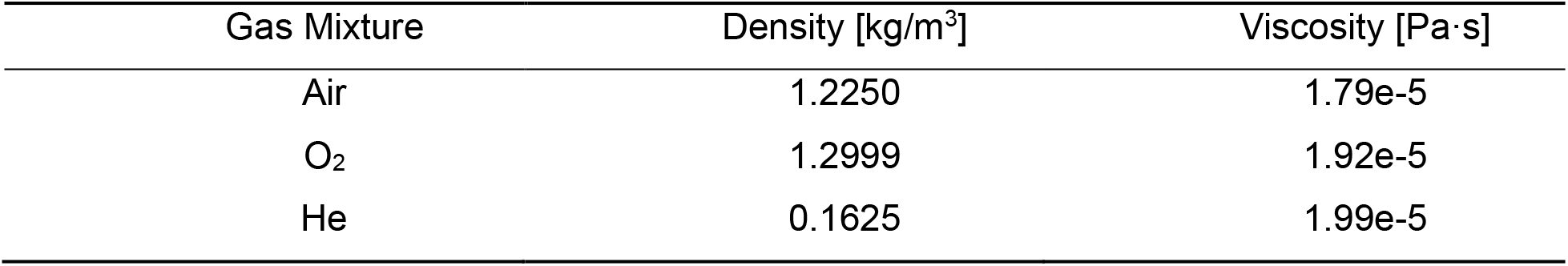
Material properties of gas species employed in this study.

| Gas Mixture | Density [kg/m<sup>3</sup>] | Viscosity [Pa·s] |
| --- | --- | --- |
| Air | 1.2250 | 1.79e-5 |
| O<sub>2</sub> | 1.2999 | 1.92e-5 |
| He | 0.1625 | 1.99e-5 |

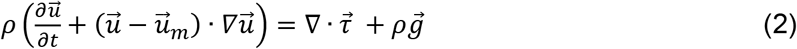

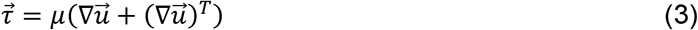

where *ρ* is the gas mixture density, *p* is pressure, 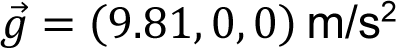 is the gravitational acceleration, 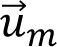 is deformable mesh velocity vector, 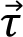 is viscous stress tensor, *μ* is the dynamic viscosity of gas mixture. In this study, the density and viscosity of gas mixtures were calculated from the volume fractions of gas constituents. Because of the airway deformation kinematics in DLAM, the convective velocity term, 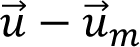, in **Eq. (2)** is induced by the difference between the localized air velocity and the dynamic mesh velocity. 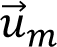 is defined by the temporal derivative of coordinates 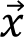 of dynamic mesh nodes, i.e.,

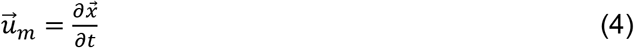

Specifically, with the no-slip boundary conditions at airway walls, gas flow velocity on moving airway boundaries 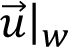 can be expressed as:

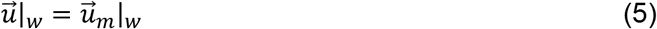

For SLAM simulations, 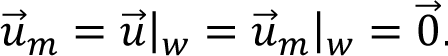.

#### 2.2.2 Discrete phase (aerosolized medicinal particles)

To investigate the particle size influence on particle transport and deposition, three different sizes of monodispersed and spherical injected particles at the mouth opening were considered (i.e., aerodynamic diameter *d_p_*=1, 5, and 10 *μm*, respectively. Accordingly, particle density *ρ_p_* = 1000 *kg*/*μ*³. Particle trajectories were predicted by solving the particle translation equation (i.e., Newton’s 2^nd^ law), where the drag force 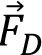, Saffman lift force 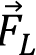, and gravity 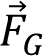 as the primary forces (Lukerchenko et al., 2012) acting on particles. Brownian motion-induced force was neglected, because it is negligible for particles with a diameter equal to or larger than 1 µm (Heyder et al., 1986). Specifically, the particle translation equation can be given as

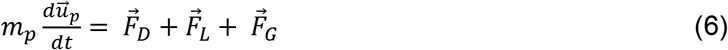

where *μ_p_* and 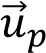 represent the particle mass and velocity, respectively. 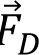 is given by

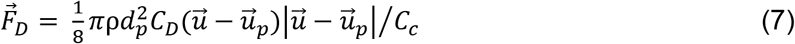

where *d_p_* denotes the particle diameter, *C_c_* represents the Cunningham correction factor (Allen & Raabe, 1985), and *C_D_* refers to the drag coefficient (Chen et al., 2018) given by

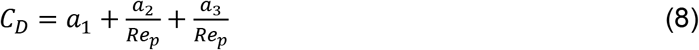

The constants *a*_1_, *a*_2_, and *a*_3_ can be determined based on the particle Reynolds number (*Re_p_*), which is defined as (Morsi & Alexander, 1972)

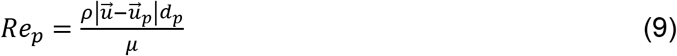

The Cunningham correction factor *C_c_* in **Eq. (7)** is defined as (Allen & Raabe, 1985)

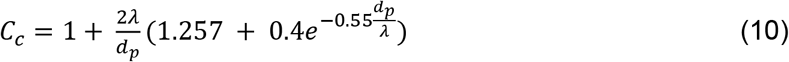

in which *λ* is the mean free path of air or heliox mixture. It is worth noting that *C_c_* was evaluated using the air-based correlation, while carrier gas dependence was incorporated through the gas-specific *λ* values. **Table 2** provides *C_c_* values of for particles with different diameters and gas carriers employed in this study. Furthermore, the tensor form of the Saffman lift force *F_L,i_* can be given as (Li & Ahmadi, 1992)

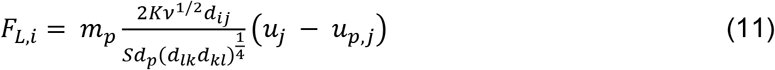

**Table 2:** Cunningham correction factor under various particle sizes and gas mixtures.

| $d_p$ [μm] | 1 | 5 | 10 |
| --- | --- | --- | --- |
| Air | 1.167 | 1.033 | 1.017 |
| He-O<sub>2</sub> | 1.265 | 1.051 | 1.026 |

where *K* = 2.594 is the constant coefficient for the Saffman lift force, *ν* represents the kinematic viscosity, *S* is the ratio of particle density to gas flow density, and *d_ij_* is the deformation rate tensor, which is defined as

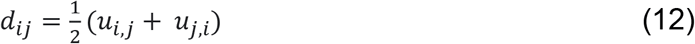

### 2.3 Glottis motion model

The dynamics of the glottis motion were modeled based on available clinical measurements (Scheinherr et al., 2015; Scheinherr et al., 2012) which were employed in our previous publication (Zhao et al., 2020). Assuming the anterior-posterior length at the cross-section of the glottis remains constant (i.e., Plane A in **Fig. 2 (a)**). In any inhalation-exhalation breathing cycle with period equal to *T* (see **Fig. 3**). Based on the fact that the displacement of each mesh node is a function of both time and space in the dynamic glottis region, a clinically validated and computationally optimized generalized glottis motion function was used in this study to mimic the glottis motion (Scheinherr et al., 2015; Scheinherr et al., 2012) with a user-defined function (UDF) (Zhao et al., 2020). Nodal displacement of the glottis motion can be expressed as

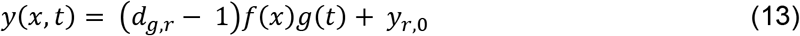

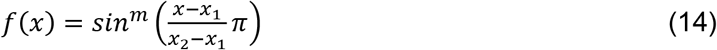

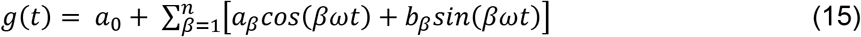

where *y_r_*_,0_ is the initial *y*-coordinate of the surface mesh nodes of the dynamic glottis region (see the red-highlighted region in the upper airway in **Fig. 1 (a)**), and *d_g,r_* is the deformation ratio between the maximum glottis width at 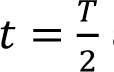 and the glottis width at the neutral position *t* = 0, which was equal to 1.4 in this study (Zhao et al., 2020). *f*(*x*) describes the heterogeneous motion of the glottal region along the *x*-direction to achieve a smooth transition from the maximum vocal fold deformation (*x* = 0.065 m) to the zero *y* displacement airways (*x* ≤ 0.053 m and *x* ≥ 0.073 m). The positive exponent can be optimized based on different intensities of glottis adduction and abduction. Accordingly, *x*_1_ and *x*_2_ are 0.053 and 0.073 m, respectively. They are determined based on the current coordinates of the mouth-to-G9 geometry. *μ* indicates the fast decay of the deformation ratio in the *x*-direction from the vocal fold to *x*_1_ and *x*_2_, which was equal to 4 in this study (Zhao et al., 2020). *g(t)* is the nodal displacement function, based on a time-dependent Fourier series function. As an example, **Fig. 2 (a)** shows aperture variations of a selected plane (i.e., Plane A) in the dynamic glottis region at multiple representative time stations associated with sinusoidal breathing waveforms (see **Fig. 3**) over one inhalation-exhalation cycle, of which the inhalation-to-exhalation duration ratio is 1:1.

### 2.4 TB tree motion model

The TB tree motion (see **Fig. 2 (b)**) is achieved by simulating anisotropic expansion and compression with realistic volume changes based on pulmonary function test (PFT) statistics (Bates, 2009; West & Luks, 2021; Xi et al., 2018). One-way coupling between airway motion and gas flow is assumed in this study. Specifically, since air and heliox have low density, the gas flow influence on airway deformation is negligible, compared with spontaneous airway motion driven by the diaphragm. Accordingly, temporal and spatial nodal displacements of the TB tree from trachea to G9 (see the lower red highlighted region in **Fig. 1 (b)**) are mimicked using dynamic mesh strategy implemented via an in-house UDF. At time *t*, the coordinate *x_i_* (*s* = 1, 2, 3) of mesh nodes in the TB tree can be calculated using

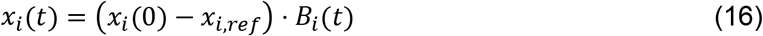

where *x_i_*(0) is the initial coordinates of mesh nodes at the start of inhalation, and *x_i,Ref_* is the coordinates of the stationary reference point, i.e., the geometric center of the reference plane (see **Fig. 2 (b)**). The transformation function *B_i_(t)* can be given as:

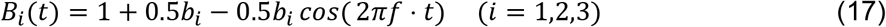

where *f* [Hz] is the frequency, i.e., the number of breath cycles per second. *b_i_* is the deformation ratio vector, of which the value varies with direction and lung health condition (Zhao et al., 2021). The anisotropic deformation ratio along the three directions was given by statistics with previous investigations (Bates, 2009; De Groote et al., 1997; Plathow et al., 2004; West & Luks, 2021; Xi et al., 2018; Zhao et al., 2021). In this study,

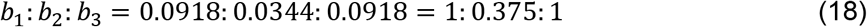

The restricted deformation ratio along the left-to-right y-direction (i.e., *b*_2_) is limited by the rib cage.

### 2.5 Two-stage formulation of inhalation-exhalation flow rate determination

Because the subject-specific computational geometry terminates at G9, deformation of the resolved mouth-to-G9 domain accounts for only part of the volume change associated with whole-lung expansion and contraction. Expansion and contraction of the resolved airway geometry (i.e., mouth-to-G9) induce airflow through the mouth opening, but this deformation-induced flow does not represent the ventilation flow rate of the unresolved distal airways and alveolar region downstream of G9. Therefore, a two-stage procedure was employed to establish a physiologically representative inhalation-exhalation waveform for the DLAM simulations and an equivalent flow condition for comparison with SLAM.

#### 2.5.1 Stage 1: Determination of the DLAM-equivalent total inspired volume

A baseline sinusoidal volumetric flow rate (VFR) waveform was first used to represent ventilation of the unresolved distal airways. The baseline VFR (*VFR_base_*) was defined as

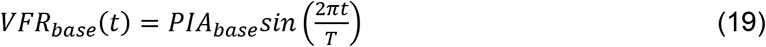

where *T* is the breathing-cycle period and *PIA_base_* is the peak inhalation flow rate. Based on physiological pulmonary function data (De Groote et al., 1997; Plathow et al., 2004; Xi et al., 2018), a target tidal volume (*TV*) of 7 mL/kg of ideal body weight was employed, corresponding to a baseline *TV_base_* of approximately 630 mL for the subject used in this study (see **Section 2.1**). Accordingly,

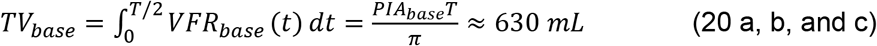

For the 12 BPM case (*T* = 5 s), this baseline waveform is shown by the solid red curve in **Fig. 3**. For Stage 1, the baseline waveform was used to define the velocity boundary conditions at the G9 terminals, while the prescribed glottis and TB tree deformation were activated in DLAM. The time-dependent expansion and contraction of the resolved mouth-to-G9 domain generated an additional deformation-induced contribution to the flow rate measured at the mouth opening. The resulting DLAM mouth VFR can be expressed as

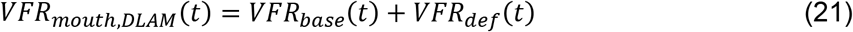

where *VFR_def_(t)* represents the flow rate contribution associated with the time-dependent volume change of the resolved mouth-to-G9 domain. The instantaneous VFR at the mouth opening was recorded throughout the breathing cycle. Integration over the inspiratory phase yielded a total DLAM inspired volume of approximately 710 mL, i.e.,

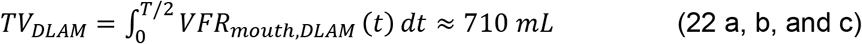

#### 2.5.2 Stage 2: Construction of an equivalent breathing waveform for SLAM-DLAM comparison

To isolate the effects of airway deformation from variations in total inhaled volume, the time-resolved *VFR_mouth,DLAM_(t)* data obtained in Stage 1 from DLAM simulation was recorded and exported as a tabulated transient profile, which is later applied to SLAM simulation. The tabulated transient profile (time vs. VFR) follows a quasi-sinusoidal waveform. At 12 BPM (*T* = 5 *ss*), the peak inhalation flow rate of *VFR_mouth,DLAM_(t)* was approximately 446 mL/s, compared with approximately 396 mL/s for the original 630-mL baseline waveform shown by the solid red curve in **Fig. 3**. The recorded DLAM *VFR_mouth,DLAM_(t)* data were then prescribed at the G9 terminal openings of the SLAM. Because the SLAM geometry is static, the imposed G9 flow directly determines the mouth flow, allowing the SLAM to reproduce the same overall mouth inhalation profile and inspired volume as the DLAM. Thus, the SLAM-DLAM comparison was performed under matched inhalation-exhalation flow rate profiles, retaining airway deformation as the principal difference between the two models. For the subsequent DLAM parametric study, the same tidal volume of 710 mL was maintained, while the breathing frequency varied to 12, 16, and 20 BPM, corresponding to breathing periods *T* = 5, 3.75, and 3 s, respectively. The associated transient *VFR_mouth,DLAM_ (t)* tabulated quasi-sinusoidal waveform profiles, other than the red solid curves, were shown in **Fig. 3**. This design allows breathing-frequency effects to be evaluated at a fixed tidal volume while preserving a consistent basis for comparison among the investigated cases.

### 2.6 Particle regional deposition and delivery fractions

To quantify regional airway aerosol dosimetry, two different metrics were employed based on different particle fates. Particles deposited on airway surfaces within the resolved mouth-to-G9 geometry were quantified using the regional deposition fraction (DF), whereas particles exiting through the G9 terminals were quantified using the beyond-G9 delivery fraction (also denoted as DF) but with different subscripts. Because the computational geometry terminates at G9, particles crossing the G9 outlets were considered delivered to the unresolved distal airways but were not treated as explicitly resolved deposition within those regions. For a specific airway region, the regional DF was calculated as

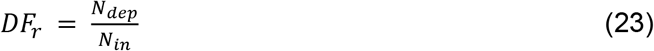

where *N_dep_* is the number of particles deposited on airway surfaces within region *r* (i.e. *r* = mouth-to-trachea or G0-G9), and *N_in_* is the total number of inhaled particles.

Since the present subject-specific airway geometry extends only to G9, deposition in the smaller bronchioles, acinar airways, and alveolar regions was not explicitly resolved. Instead, particles escaping G9 terminals were assumed to remain in the downstream respiratory tract and were used as a measure of distal aerosol delivery. Accordingly, the beyond-G9 delivery fraction was defined as

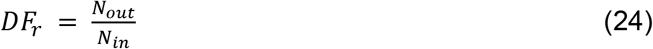

where subscript *r* (i.e., *r* = beyond-G9), and *N_out_* is the number of particles exiting through the G9 terminals. Under the present modeling assumption, particles exiting the G9 terminals were considered retained in the downstream airways and cannot re-enter the mouth-to-G9 computational domain.

At the end of simulations, a small fraction of particles remained suspended within the mouth-to-G9 computational domain. These particles were not assigned either to a regional deposition fraction or to the beyond-G9 delivery fraction. Instead, they were quantified using the residual suspended particle fraction (*R_susp_*), which is defined as

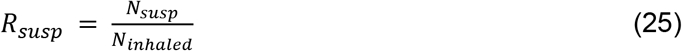

where *N_susp_* is the number of particles remaining suspended within the mouth-to-G9 computational domain at the end of inspiration. **Table 3** presents *R_susp_* values for all simulation cases in this study. It is worth noting that *R_susp_* was used only as a reference when interpreting very small potential variations in regional deposition fractions and beyond-G9 delivery fractions.

**Table 3:** Residual suspended-particle fractions (*R_susp_*) for SLAM and DLAM simulations with different carrier gases, breathing frequencies, and particle diameters.

| Lung Type | Gas | BPM | $d_p$ [μm] | | |
| --- | --- | --- | --- | --- | --- |
|  |  |  | 1 | 5 | 10 |
| | | | $R_{susp}$ | | |
| Rigid | Air | 12 | 0.22% | 0.09% | 0.87% |
| Rigid | Heliox | 12 | 0.30% | 0.20% | 0.82% |
| DLAM | Air | 12 | 0.33% | 1.08% | 1.11% |
| DLAM | Heliox | 12 | 0.42% | 0.21% | 0.78% |
| DLAM | Air | 16 | 0.22% | 3.09% | 1.62% |
| DLAM | Heliox | 16 | 0.37% | 0.26% | 0.98% |
| DLAM | Air | 20 | 0.23% | 0.25% | 1.40% |
| DLAM | Heliox | 20 | 0.31% | 0.52% | 1.41% |

### 2.7 Numerical setup

#### 2.7.1 Continuous phase

To quantify the dynamic lung effects (i.e., glottis and TB tree motions) on flow characteristics between air and heliox in airways and associated medicinal particle transport and deposition, this study used identical inhalation-exhalation boundary conditions (BCs) with the inhalation-exhalation cycle period *T* = 5 *s* (i.e., BPM = 12) for both SLAM and DLAM. A realistic quasi-sinusoidal waveform (see red dashed waveform in **Fig. 3**) was employed in both SLAM and DLAM, which was derived from the baseline BC as discussed in **Section 2.5**.

Furthermore, to investigate the generality of heliox influence on inhaled particles delivery into small airways, i.e., downstream to G9, three inhalation-exhalation sinusoidal waveforms were applied at the small airway terminals (see **Fig. 3**) with three breathing frequencies (i.e., 12, 16, and 20 BPM), a fixed tidal volume of 710 ml. The inhalation and exhalation time duration ratio was set as 1:1. For each CFPD simulation in this study, three inhalation-exhalation cycles were simulated under the three designated BPMs mentioned above, with the corresponding periods (i.e., *T* = 5, 3.75, and 3 *s*), respectively. Additionally, in each CFPD simulation, particles were injected at the start of the third inhalation-exhalation cycle. Except for the glottis region and distal trachea-G9, other airway walls were assumed stationary and non-slip in this study. Outlet vent BC was assigned at the mouth opening, with a gauge pressure equal to 0 Pa.

#### 2.7.2 Discrete phase

Particles were injected into the mouth opening at the beginning of inhalation-exhalation cycle and released four times. The first release was at a physical flow time of 0.005 s. The other three releases proceeded with a time interval of 0.005 s. A total of 100,000 particles were injected for each simulation case, and a single release was embedded with 25,000 particles. The time step and total number of particles were determined based on time-step and particle number independence tests. It is worth mentioning that at the mouth opening, particles can escape during exhalation. Particles were designated to be trapped at the airway walls. It means once the distance between the centers of the particles and the wall is less than the particle radius, they are considered deposited. At the G9 terminals, escaped BCs were assigned for particles. Particles that exit from G9 terminals were assumed to be deposited 100% in G9-to-alveoli regions, i.e., no particles can reenter the mouth-to-G9 domain after escaping.

#### 2.7.3 Numerical implementation and solver settings

CFPD simulations were performed using Ansys Fluent 2025 R2 (Ansys Inc., Canonsburg, PA) with 48 parallel solver processors on a local Dell T7960 workstation (Intel® Xeon® Processor W9-3575X with dual processors, 44 cores, 88 threads, and 256 GB RAM), in which, 16 cores with 32 threads were used in the current simulations. With the flow time step equal to 0.005 s, the simulations required approximately 28, 36, and 45 hours to complete DLAM simulations for half of an inhalation-exhalation cycle with BPM = 20, 16, and 12, respectively. For SLAM simulations with *T* = 5 s, it took approximately 35 hours for one case simulation, less than the corresponding DLAM case.

Finite Volume Method (FVM) with a Semi-Implicit Method for Pressure Linked Equations (SIMPLE) scheme was employed for solving the governing equations. The least squares cell-based (LSCB) method was employed to calculate cell-average gradients. In addition, a second-order scheme was used for pressure discretization. A second-order upwind approach was used for the discretization of momentum, turbulence kinetic energy, and specific dissipation rate in both time and space. It is considered that the simulations have converged when all residuals are below 1 × 10^-3^. In-house UDFs and MATLAB scripts were employed for solver customization to achieve the following capabilities, i.e.,

1. Defining the prescribed deformation kinematics of the glottis and distal trachea-G9 configurations with different breathing patterns,
2. Generating particle injection files,
3. Defining the transient sinusoidal inhalation-exhalation waveforms,
4. Specifying the DPM time step, and
5. Post-processing particle deposition data as well as WSS on the airway wall.

### 2.8 Model validations and calibrations

#### 2.8.1 Euler-Lagrange model validation

The Euler-Lagrange model employed in this study was validated by comparisons with benchmark experimental data on pulmonary airflow field and regional particle deposition. Details of the validations can be found in previous publications (Feng et al., 2015; Feng et al., 2017; Haghnegahdar et al., 2019; Yi & Feng, 2026). Specifically, the flow fields were resolved using the transition SST model, which is selected based on prior validations (Chen et al., 2017; Feng et al., 2017; Jiang et al., 2025; Li et al., 2025; Liu et al., 2024; Tan et al., 2008; Zhang & Kleinstreuer, 2011) demonstrating its accuracy in predicting laminar-to-turbulence transitional pulmonary gas flow fields and particle deposition.

#### 2.8.2 Dynamic mesh calibration and validation

Previous investigations established and evaluated the in-house dynamic mesh method used in this study to prescribe glottal motion and TB tree deformation kinematics (Zhao et al., 2020; Zhao et al., 2021). For the glottal motion model described by **Eqs. (13)-(15)**, Zhao et al. (2020) calibrated the prescribed glottal abduction and adduction using clinical imaging measurements. The resulting time-dependent glottis aperture agreed well with clinical data under both idealized sinusoidal and clinically measured breathing waveforms. Furthermore, the resulting flow dynamics, especially the variation trends in average mainstream pressure drops induced by the dynamic mesh, were qualitatively validated against experimental aerodynamic data derived from physical airway casts.

The prescribed expansion and contraction of the TB tree (see **Eqs. (16)-(18)**) are derived from the deformable truncated whole-lung framework developed by Zhao et al. (2021). In that study, generalized airway deformation functions were calibrated using clinically reported lung volume changes for both healthy subjects and patients with COPD. The predictions of lung-volume variation throughout the respiratory cycle demonstrated good agreement with available physiological and experimental data. These previously calibrated deformation relationships provide the physiological foundation for prescribing the time-dependent lobar and airway motion used in the present DLAM.

## 3. Results and Discussion

This section examines two related research questions. The first question is on the extent to which physiologically prescribed airway deformation kinematics affect gas-particle flow dynamics and regional aerosol dosimetry compared with the conventional SLAM. To address this, **Section 3.1** directly compares SLAM and DLAM at 12 BPM, using an identical inhalation-exhalation waveform for both air and heliox and considering particle sizes of 1, 5, and 10 µm. The second question investigates how carrier gas properties and breathing frequency influence airflow characteristics, regional particle deposition within the resolved mouth-to-G9 airways, and particle delivery beyond G9, after accounting for the effects of airway deformation. Therefore, **Section 3.2** employs DLAM to compare air and heliox at 12, 16, and 20 BPM, maintaining a constant tidal volume of 710 mL for all three particle sizes.

### 3.1 SLAM vs. DLAM

#### 3.1.1 Influence on gas flow field

**Figure 4** illustrates a direct comparison of the transient velocity fields along the sagittal plane at Y=0.0034 m (i.e., Plane B in **Fig. 4 (a)**) and glottal cross-section X=0.06405 m (i.e. Plane C in **Fig. 4 (a)**). The same equivalent inhalation-exhalation waveform (see dashed red curve in **Figs. 3 and 4(a)**) was employed for this comparison. Velocity fields predicted using SLAM and DLAM are presented at three representative inhalation time stations (i.e., *t* = *T*/8, *T*/4, and 3*T*/8), respectively. On Plane B, SLAM predicts local maximum velocity magnitudes 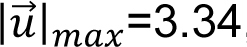, 4.88, and 3.68 m/s at the three respective time points, whereas the corresponding DLAM values are 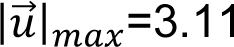, 4.52, and 3.43 m/s, as shown in **Figs. 4 (b) and (c)**. Thus, relative to DLAM, SLAM predicts higher localized velocity magnitudes along the sagittal plane (i.e., Plane B) throughout the investigated inhalation phase, with the largest absolute difference occurring near peak inspiration at *t* = *T*/4. On Plane C, with the larger surface areas at all time stations due to the abduction (i.e., expansion) of the glottis during the inhalation, DLAM predicted lower localized velocity magnitudes than SLAM at the glottis (i.e., Plane C). This behavior is associated with the prescribed glottal abduction and TB tree expansion in DLAM, which continuously enlarges the local cross-sectional area during inhalation and introduces dynamic mesh motion into the gas-flow field. In contrast, the SLAM maintains a fixed airway geometry throughout the breathing cycle.

**Fig. 4.**
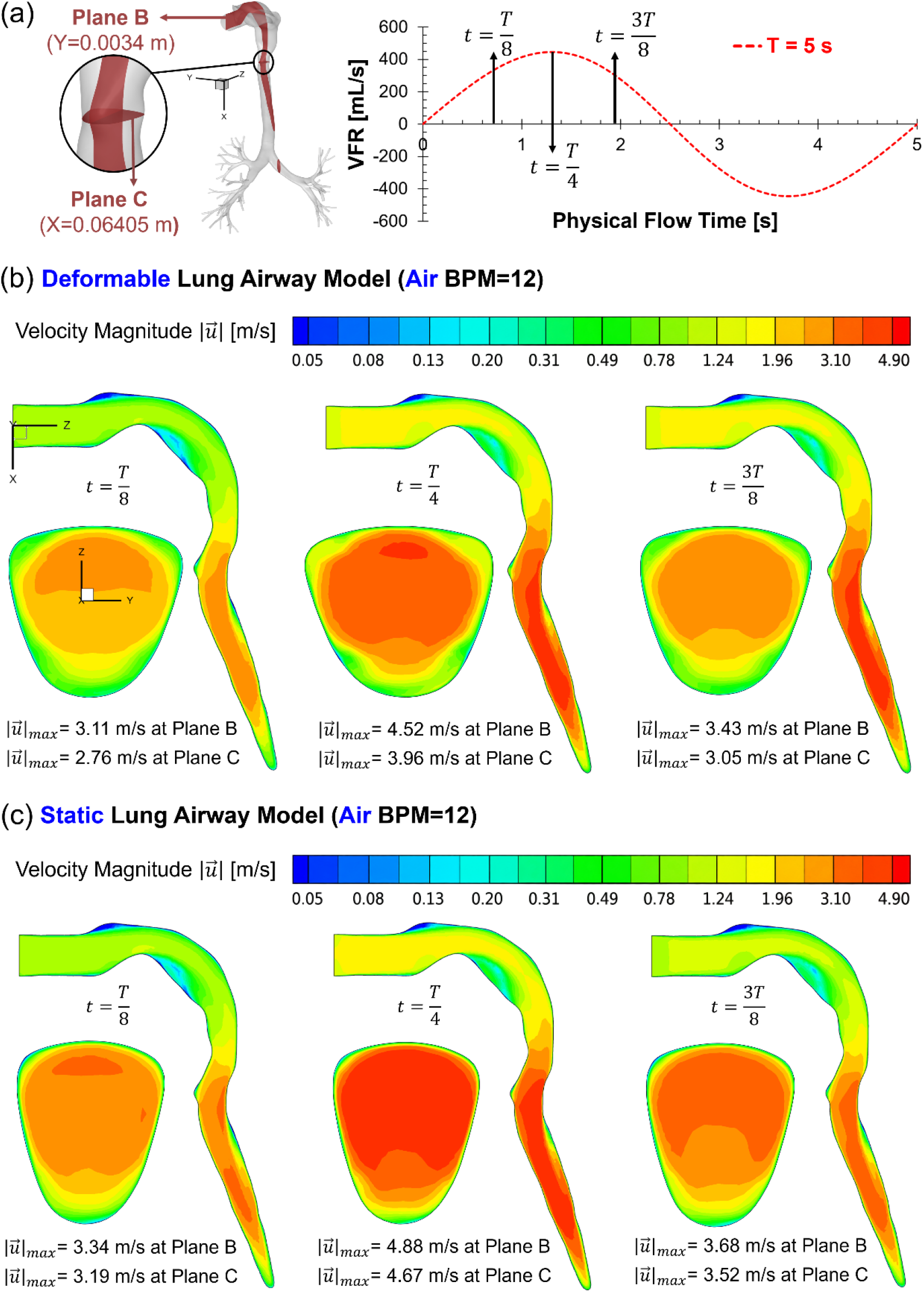
Comparison of air velocity distributions between DLAM and SLAM at 12 BPM: (a) locations of Planes B and C and the selected inhalation time points; (b) velocity magnitude distributions predicted by the DLAM; and (c) velocity magnitude distributions predicted by the SLAM.

A consistent deformation effect is also observed for heliox, as shown in **Fig. 5**, in which the SLAM and DLAM simulations also employed the identical breathing waveform (see **Fig. 5 (a)**) used for the simulations shown in **Fig. 4**. For heliox, SLAM predicts maximum velocity magnitudes 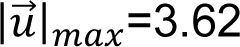, 5.15, and 3.98 m/s, whereas DLAM predicts 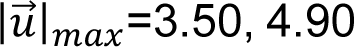, 4.90, and 3.74 m/s at *t* = *T*/8, *T*/4, and 3*T*/8, respectively (see **Figs. 5 (b) and (c)**). The differences are again more noticeable at Plane C, where the corresponding SLAM 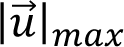 values of 3.35, 4.88, and 3.69 m/s decrease to 2.91, 4.16, and 3.21 m/s in DLAM (see **Figs. 5 (b) and (c)**). Therefore, when physiologically realistic airway deformation kinematics are neglected, the SLAM consistently overpredicts the localized velocity magnitude during inhalation relative to the DLAM. This discrepancy is noticeable especially in the glottis region, where the SLAM predicts approximately 15-18% higher maximum velocity magnitudes than the DLAM for both air and heliox at the investigated time stations during the inhalation period.

**Fig. 5.**
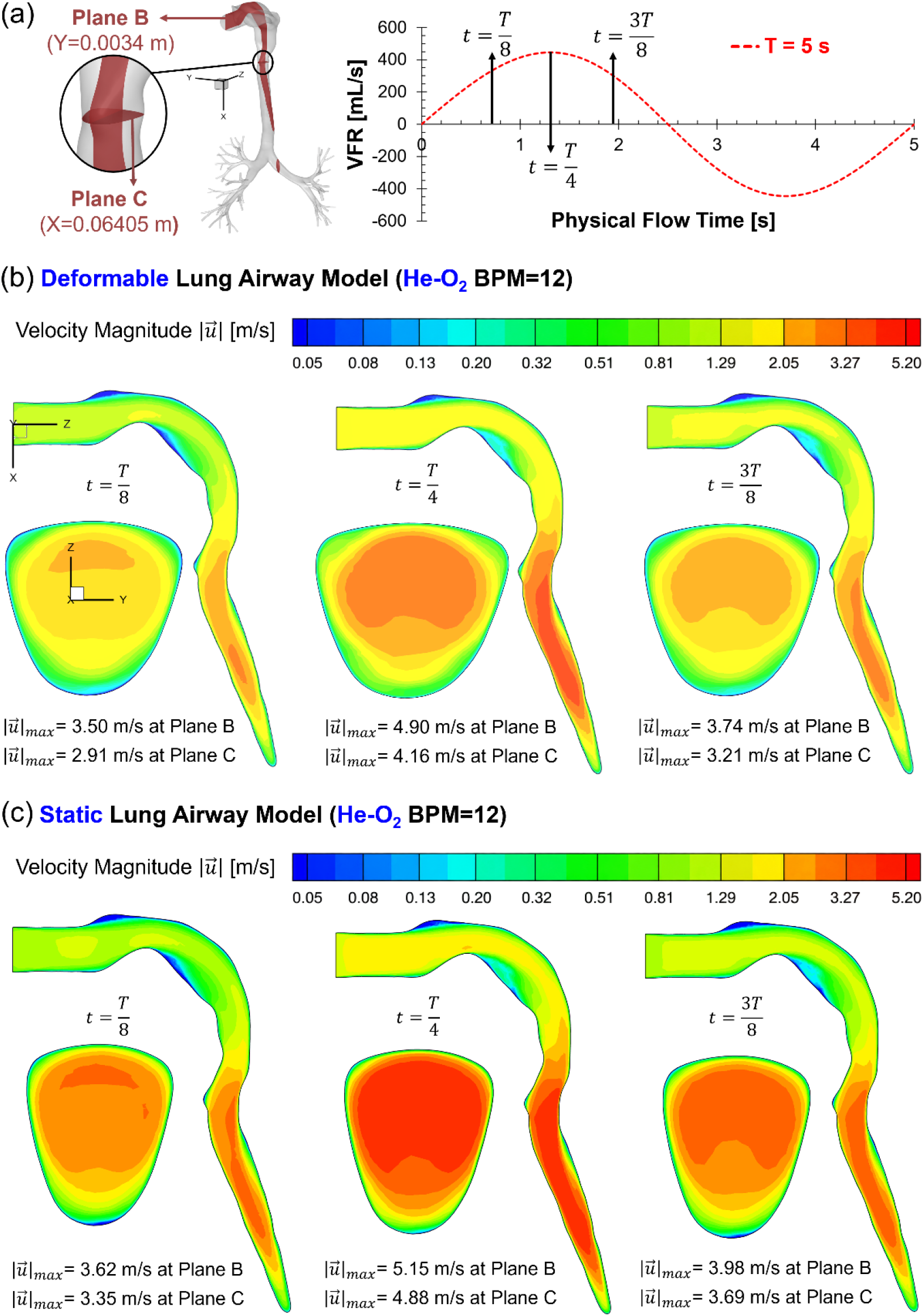
Comparison of heliox (70% He-30% O_2_) velocity distributions between DLAM and SLAM at 12 BPM: (a) locations of Planes B and C and the selected inhalation time points; (b) velocity magnitude distributions predicted by DLAM; and (c) velocity magnitude distributions predicted by SLAM.

Furthermore, as shown in **Figs. 6 and 7**, airway deformation kinematics also noticeably influence the prediction of turbulence kinetic energy (TKE). As shown in **Fig. 6** on Plane B, the DLAM predicts lower local maximum TKE (*TKE*_max_), than the SLAM at all three selected inhalation time stations. For air, *TKE*_max_ decreases from 0.07, 0.42, and 0.18 m²/s² in the SLAM to 0.03, 0.23, and 0.10 m²/s² in the DLAM at *t* = T/8, T/4, and 3T/8, respectively. These changes correspond to reductions of approximately 57%, 45%, and 44%, indicating that neglecting airway deformation can substantially overpredict localized turbulence. The deformation effect is spatially nonuniform near the glottis, i.e., on Plane C. **Figure 6** shows that the air *TKE*_max_ values predicted by the DLAM are 0.005, 0.047, and 0.014 m²/s², compared with 0.006, 0.027, and 0.016 m²/s² in the SLAM. Therefore, it is interesting that although the DLAM predicts lower TKE during early and late inhalation, it produces a relatively higher localized *TKE*_max_ at peak inhalation, as shown in **Fig. 6 (a)**. This result indicates that glottal deformation does not simply reduce turbulence uniformly. Instead, the varying glottis aperture during the inhalation creates more disturbance in the airflow, thereby inducing higher turbulence intensity near the peak inhalation time point (i.e., *t* = T/4).

**Fig. 6.**
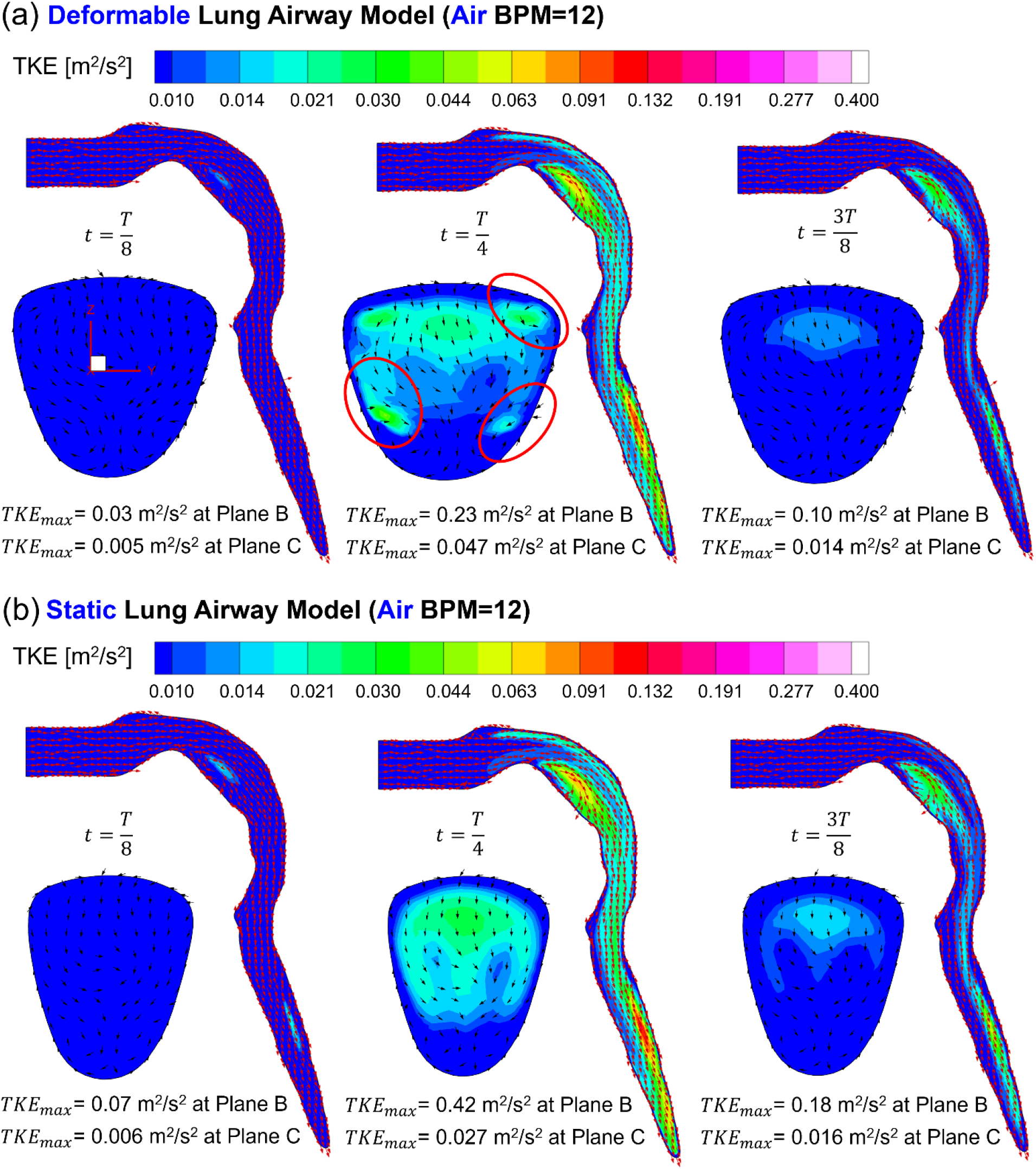
Comparison of turbulence kinetic energy (TKE) distributions and tangential (in-plane) velocity vectors of air between DLAM and SLAM at 12 BPM on Planes B and C at the selected inhalation time points: (a) DLAM and (b) SLAM

**Fig. 7.**
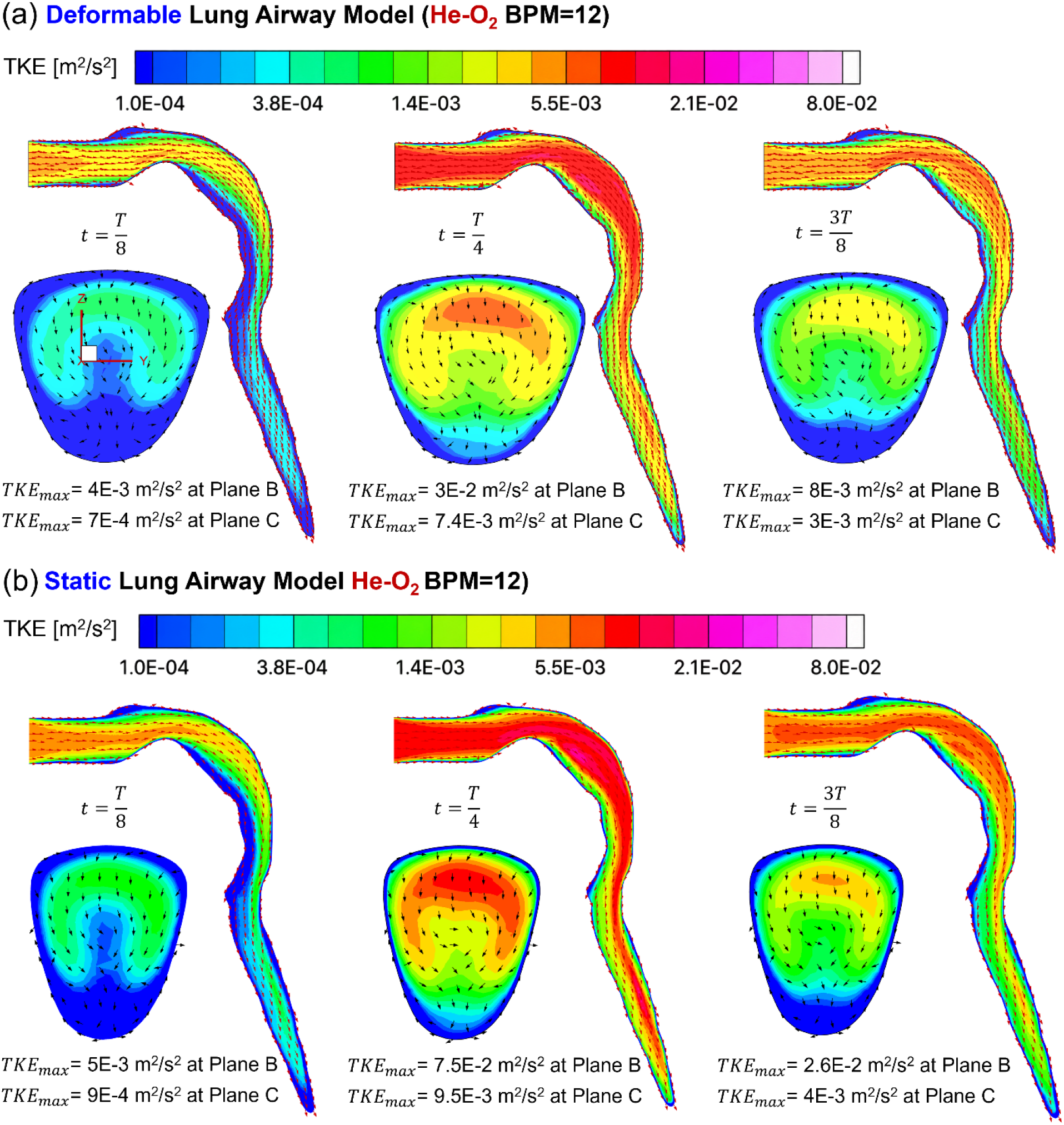
Comparison of turbulence kinetic energy (TKE) distributions and tangential (in-plane) velocity vectors of heliox (70% He-30% O_2_) between DLAM and SLAM at 12 BPM on Planes B and C at the selected inhalation time points: (a) DLAM and (b) SLAM.

A more consistent reduction for TKE is observed for heliox in **Figs. 7 (a) & (b)**. On Plane B, *TKE*_max_ decreases from 0.005, 0.075, and 0.026 m²/s² in the SLAM to 0.004, 0.030, and 0.008 m²/s² in the DLAM. On Plane C, the corresponding values decrease from 0.0009, 0.0095, and 0.004 m²/s² to 0.0007, 0.0074, and 0.003 m²/s². Therefore, for both carrier gases, incorporating physiologically realistic airway deformation kinematics noticeably changes the predicted transient turbulence field, although the magnitude and direction of the local effect depend on the airway location and inhalation phase. Together with the velocity magnitude comparisons shown in **Figs. 4 and 5**, these differences support the use of the DLAM for the subsequent evaluation of carrier gas and breathing frequency effects.

#### 3.1.2 Influence on particle transport and deposition

The gas flow field differences between SLAM and DLAM described in **Section 3.1.1** consequently influence inhaled particle transport and deposition in the human respiratory system. **Figures 8 and 9** compare the localized deposition patterns of 1, 5, and 10 µm particles with air and heliox at 12 BPM, respectively. Deposited particles are colored by their particle residence times (PRTs). The corresponding beyond-G9 delivery fractions and regional deposition fractions in the mouth-to-trachea and G0-G9 regions are presented in **Figs. 10-12**.

**Fig. 8.**
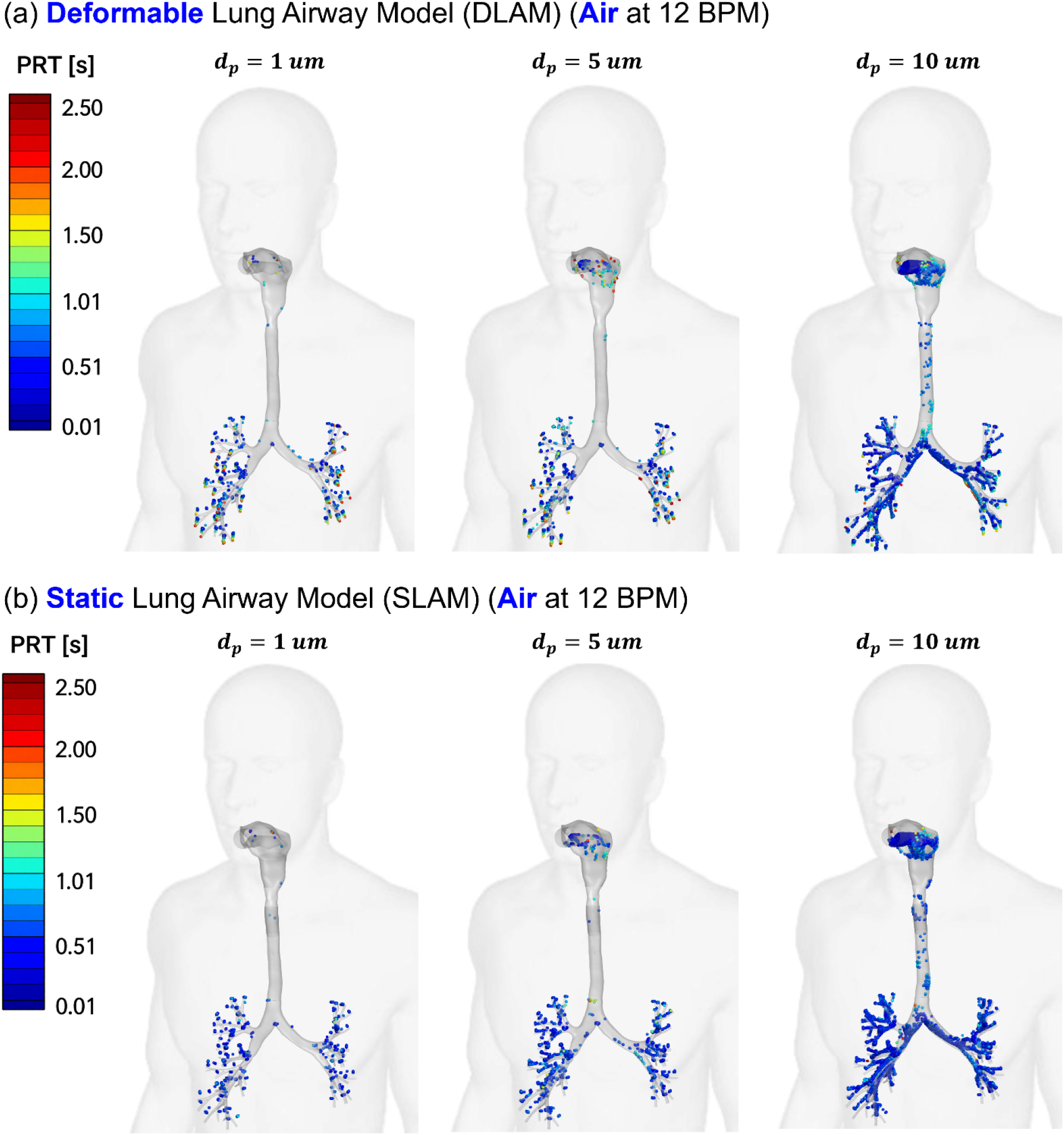
Comparison of localized particle deposition patterns between DLAM and SLAM for air at 12 BPM for particle diameters *d*_p_=1, 5, and 10 µm: (a) DLAM and (b) SLAM. Deposited particles are colored by particle residence time (PRT).

**Fig. 9.**
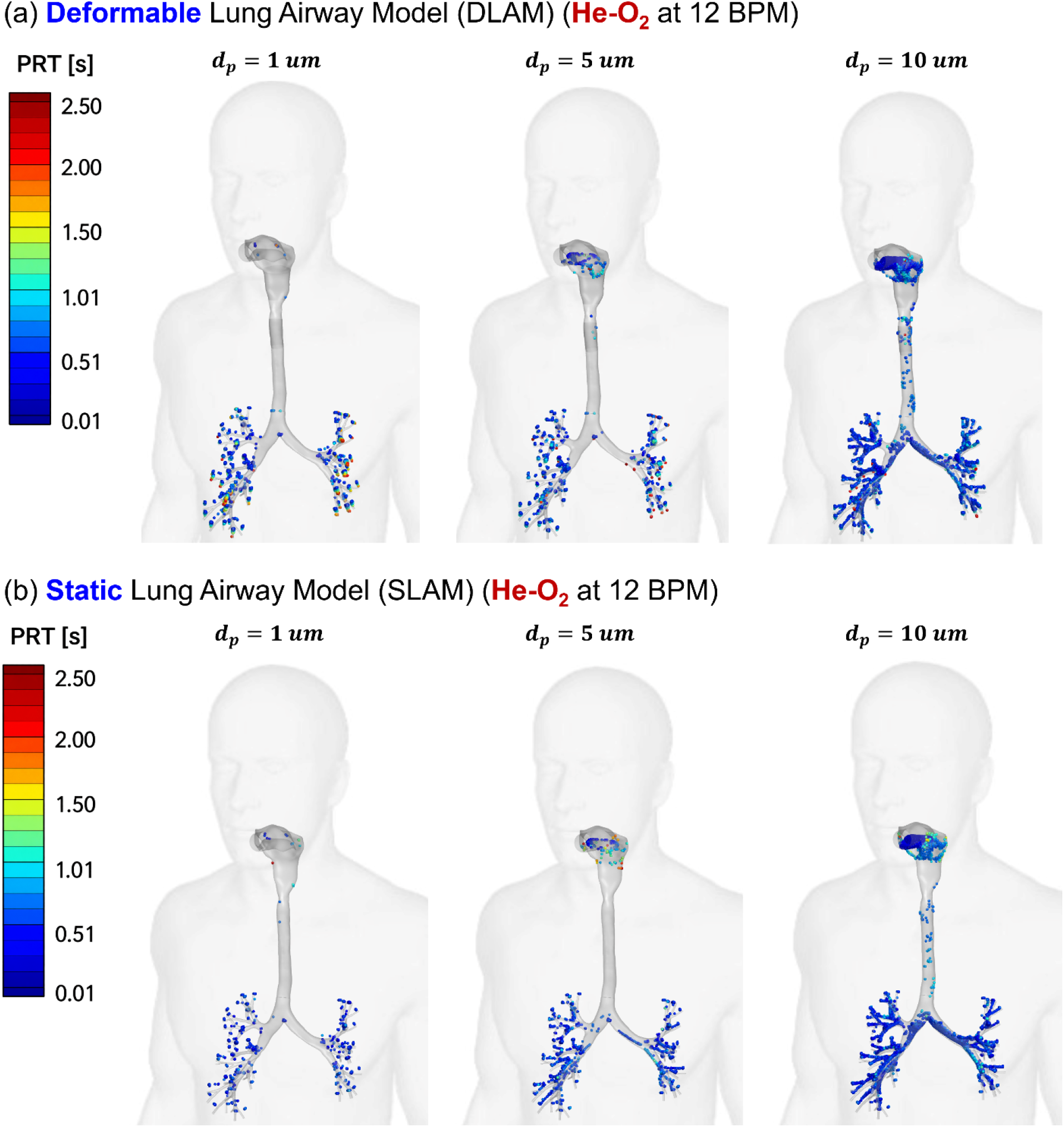
Comparison of localized particle deposition patterns between DLAM and SLAM for heliox (70% He-30% O_2_) at 12 BPM for particle diameters *d*_p_=1, 5, and 10 µm: (a) DLAM and (b) SLAM. Deposited particles are colored by PRT.

**Fig. 10.**
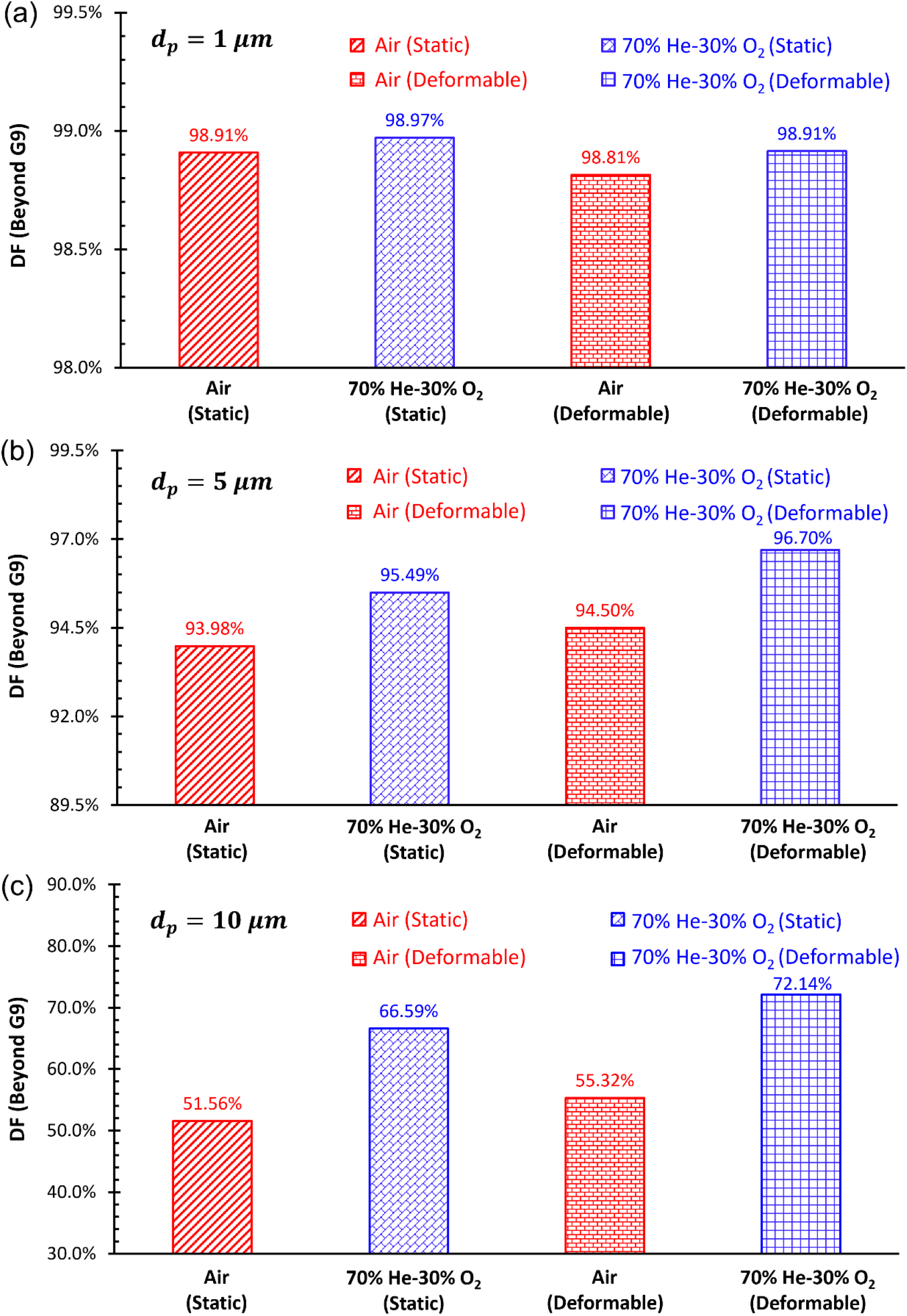
Comparison of beyond-G9 delivery fractions between SLAM and DLAM for air and heliox at 12 BPM: (a) *d*_p_=1 µm, (b) *d*_p_=5 µm, and (c) *d_p_*=10 µm.

For 1 µm particles, the influence of airway deformation is not noticeable (see **Figs. 10 (a), 11(a), and 12 (a)**). The beyond-G9 delivery fraction changes from 98.91% in the SLAM to 98.81% in the DLAM for air and from 98.97% to 98.91% for heliox (see **Fig. 10 (a)**). These absolute differences of only 0.10% and 0.06% are comparable with the residual suspended particle fractions (*R_susp_*) documented in **Table 3**. Therefore, no significant deformation-induced difference in beyond-G9 delivery can be identified for 1 µm particles under the investigated condition. Their low inertia allows them to follow the transient carrier gas flow, limiting deposition caused by either inertial impaction, gravitational sedimentation, or interception, in the region from mouth to G9.

For 5 µm particles, a clearer but still modest redistribution is observed. With air, the beyond-G9 delivery fraction increases from 93.98% using SLAM to 94.50% using DLAM (see **Fig. 10 (b)**), while the mouth-to-trachea DF increases slightly from 0.18% to 0.20% (see **Fig. 11 (b)**) and G0-G9 DF decreases from 5.75% to 4.22% (see **Fig. 12 (b)**). With heliox, the beyond-G9 delivery fraction increases from 95.49% using SLAM to 96.70% using DLAM (see **Fig. 10 (b)**), while the mouth-to-trachea DF increases from 0.19% to 0.21% (see **Fig. 11 (b)**) and G0-G9 DF decreases from 4.12% to 2.88% (see **Fig. 12 (b)**). Thus, the primary deformation-induced change occurs within the branching G0-G9 region rather than in the proximal airway. Because the absolute differences for 5 µm particles remain relatively small, particularly for air, they should be interpreted together with the corresponding *R*_susp_ values in **Table 3**.

**Fig. 11.**
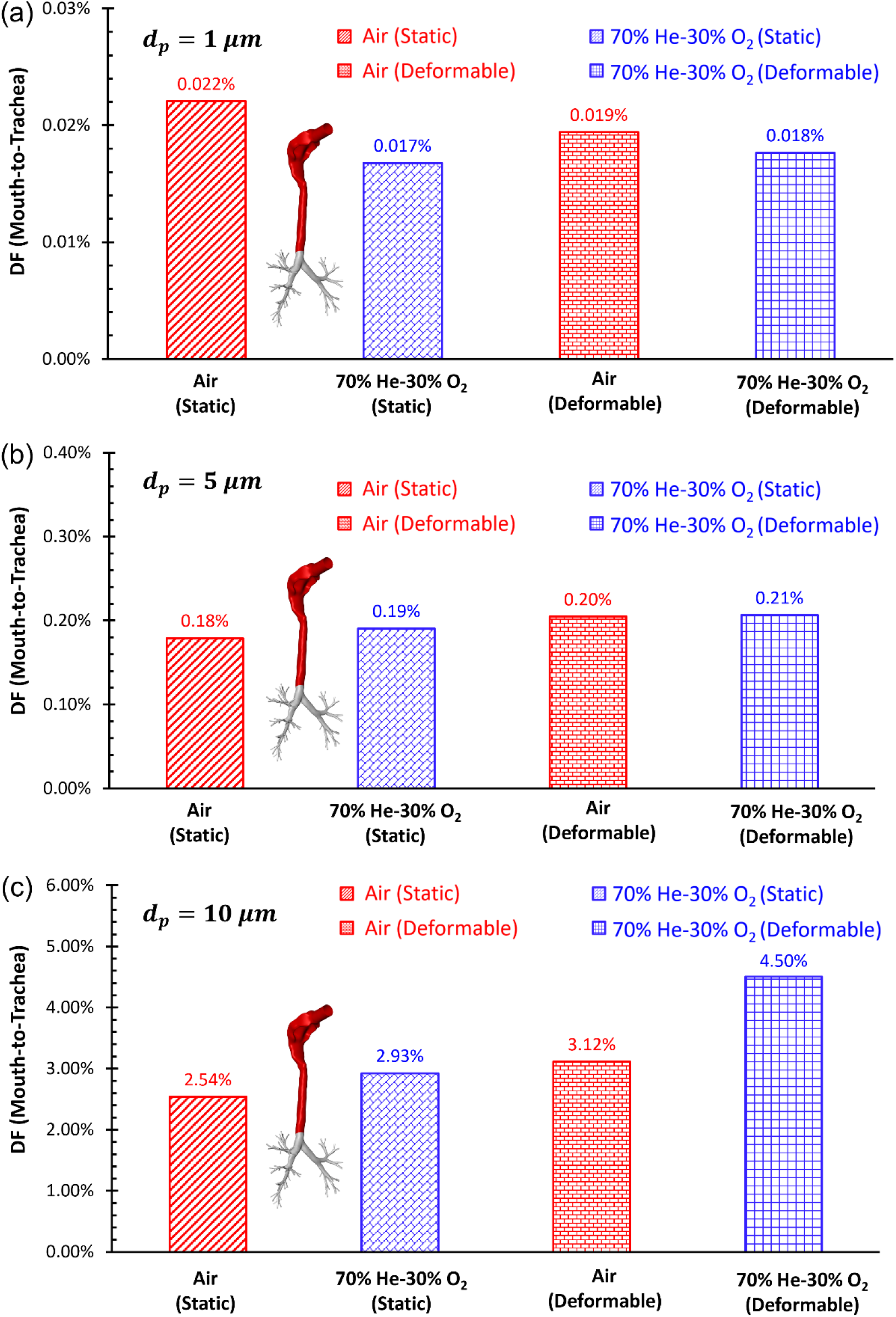
Comparison of mouth-to-trachea regional deposition fractions between SLAM and DLAM for air and heliox at 12 BPM: (a) *d_p_*=1 µm, (b) *d_p_*=5 µm, and (c) *d_p_*=10 µm.

**Fig. 12.**
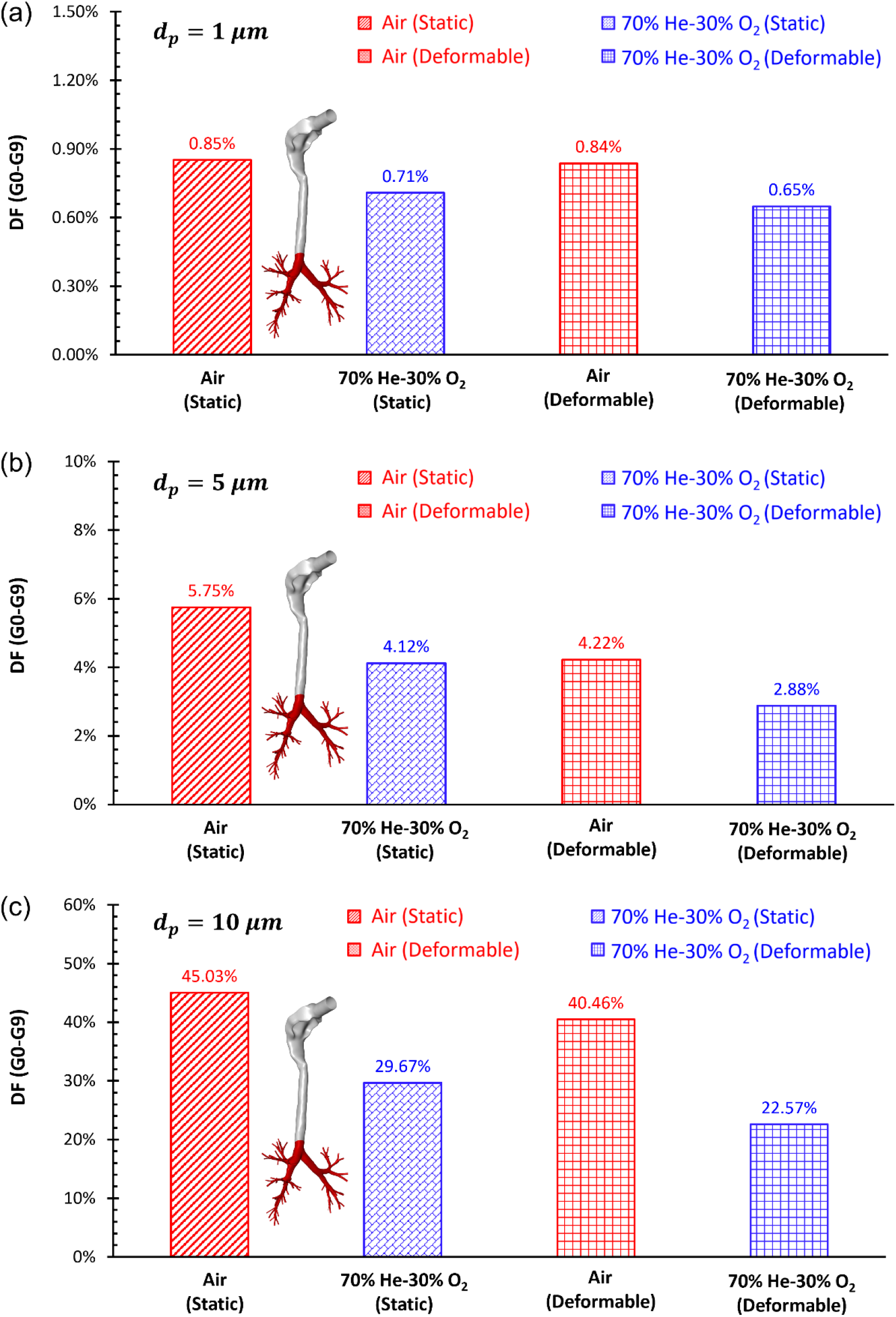
Comparison of G0-G9 regional deposition fractions between SLAM and DLAM for air and heliox at 12 BPM: (a) *d_p_*=1 µm, (b) *d_p_*=5 µm, and (c) *d_p_*=10 µm.

The influence of airway deformation kinematics becomes much more significant for 10 µm particles, as shown in **Figs. 10 (c), 11 (c), and 12 (c)**. With air, the beyond-G9 delivery fraction increases from 51.56% using SLAM to 55.32% using DLAM, an absolute increase of 3.76% (see **Fig. 10 (c)**). At the same time, the mouth-to-trachea DF increases from 2.54% to 3.12% (see **Fig. 11 (c)**) and G0-G9 DF decreases from 45.03% to 40.46% (see **Fig. 12 (c)**). With heliox, the beyond-G9 delivery fraction increases from 66.59% to 72.14% as shown in **Fig. 10 (c)**, with an absolute increase of 5.55%, while the mouth-to-trachea DF increases from2.93% to 4.50% (see **Fig. 11 (c)**) and G0-G9 DF decreases from 29.67% to 22.57% (see **Fig. 12 (c)**). These differences are much higher than the corresponding residual suspended-particle fractions shown in **Table 3**, thereby representing a clear deformation-induced redistribution of the particles.

The influence of airway deformation is highly dependent on particle size. The difference between the SLAM and DLAM is negligible for 1 µm particles, modest for 5 µm particles, and significant for 10 µm particles. For both 5 and 10 µm particles, SLAM underpredicts delivery to the beyond-G9 region and overpredicts deposition in the G0-G9 region compared with DLAM, for both air and heliox. Mouth-to-trachea deposition is higher in DLAM, with the difference being minimal for 5 µm particles and more evident for 10 µm particles. These findings suggest that airway deformation alters the transport trajectories of higher-inertia particles and redistributes their regional deposition. This redistribution results from time-dependent modifications of airway geometry and the associated gas-flow field in DLAM. In contrast to SLAM, DLAM continuously changes the local airway cross-sectional area, thereby modifying the velocity and turbulence experienced by the particles. As demonstrated in **Figs. 4-7** and further quantified by the area-averaged velocity magnitudes and TKEs at 8 representative cross-sections from the mouth to G2 in **Appendix A**, DLAM generally produces lower area-averaged velocity magnitude and TKE at most sampled locations and inhalation time points. However, these effects are spatially and temporally nonuniform, and localized increases in TKE may occur. Thus, airway deformation should be interpreted as redistributing the transient gas flow and turbulence rather than uniformly suppressing them. In airway regions where DLAM reduces local velocity magnitude and turbulent fluctuations, larger particles exhibit less inertia-driven deviation from the gas flow streamlines and reduced turbulence dispersion, resulting in less particle deposition and decreased inertial impaction. This outcome aligns with the lower G0-G9 DFs predicted by DLAM for 5 and 10 µm particles. Additionally, glottal abduction and adduction alter the local aperture, glottal jet direction, and streamline curvature. Because higher-inertia particles cannot immediately adjust to these transient flow variations, their trajectories may be redirected toward the airway wall, contributing to the higher mouth-to-trachea deposition observed with DLAM. Since the reduction in the G0-G9 deposition exceeds the corresponding increase in mouth-to-trachea deposition, a greater fraction of particles remains available for downstream transport into the beyond-G9 region. These results underscore the importance of incorporating physiologically realistic airway deformation when using CFPD to predict regional dosimetry and delivery beyond G9 for higher-inertia medicinal aerosols.

### 3.2 Gas carriers: air vs. heliox

#### 3.2.1 Influence of carriers on gas flow characteristics

After confirming the importance of airway deformation kinematics for predicting pulmonary gas-flow dynamics accurately in **Section 3.1**, DLAM was used to evaluate the effects of carrier gas and breathing frequency. Since identical breathing waveforms and airway deformation kinematics were prescribed for air and heliox (also denoted as He-O_2_), variations between the two sets of CFPD cases reflect the influence of gas carrier properties. Under identical volumetric-flow waveforms and prescribed airway deformation kinematics, the much lower density and slightly higher viscosity of heliox than air reduces the local Reynolds number and suppresses turbulence generation and inertial pressure losses. As a result, heliox produces lower TKE, WSS, and mouth-to-G9 pressure drop than air, despite comparable or slightly higher localized velocity maxima.

As an example, **Fig. 13** compares the distribution of transient velocity magnitude 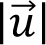 on Plane B at *t* = T/8, T/4, and 3T/8 using DLAM between air and heliox with three inhalation-exhalation waveforms. Air and heliox exhibit similar overall spatial flow patterns, while the localized maximum velocity magnitude 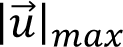 increases with breathing frequency for both gases. Heliox produces moderately higher local maxima than air. At 12 BPM (see **Fig. 13 (b)**), 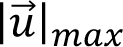 increase from 3.11, 4.52, and 3.43 m/s with air to 3.50, 4.90, and 3.74 m/s with heliox at the three selected time points, respectively. At peak inhalation (*t* = T/4), the corresponding values increase from 5.96 to 6.31 m/s at 16 BPM (see **Fig. 13 (c)**) and from 7.42 to 7.70 m/s at 20 BPM (**Fig. 13 (d)**). Thus, the use of heliox does not necessarily reduce localized velocity magnitude, even though its lower density noticeably lower TKE than air (see **Fig. 14**). Specifically, as shown in **Figs. 14 (a)-(c)**, heliox produces lower *TKE*_max_ than air at all three BPMs and time stations. At peak inhalation (*t* = T/4), *TKE*_max_ decreases from 0.24 to 0.03 m²/s² at 12 BPM, from 0.55 to 0.09 m²/s² at 16 BPM, and from 0.84 to 0.24 m²/s² at 20 BPM. These changes correspond to reductions of approximately 87.5%, 83.6%, and 71.4%, respectively. Therefore, despite the slightly higher localized velocity maxima, heliox substantially attenuates turbulence generation and yields markedly lower TKE levels in the pulmonary flow field compared with air.

**Fig. 13.**
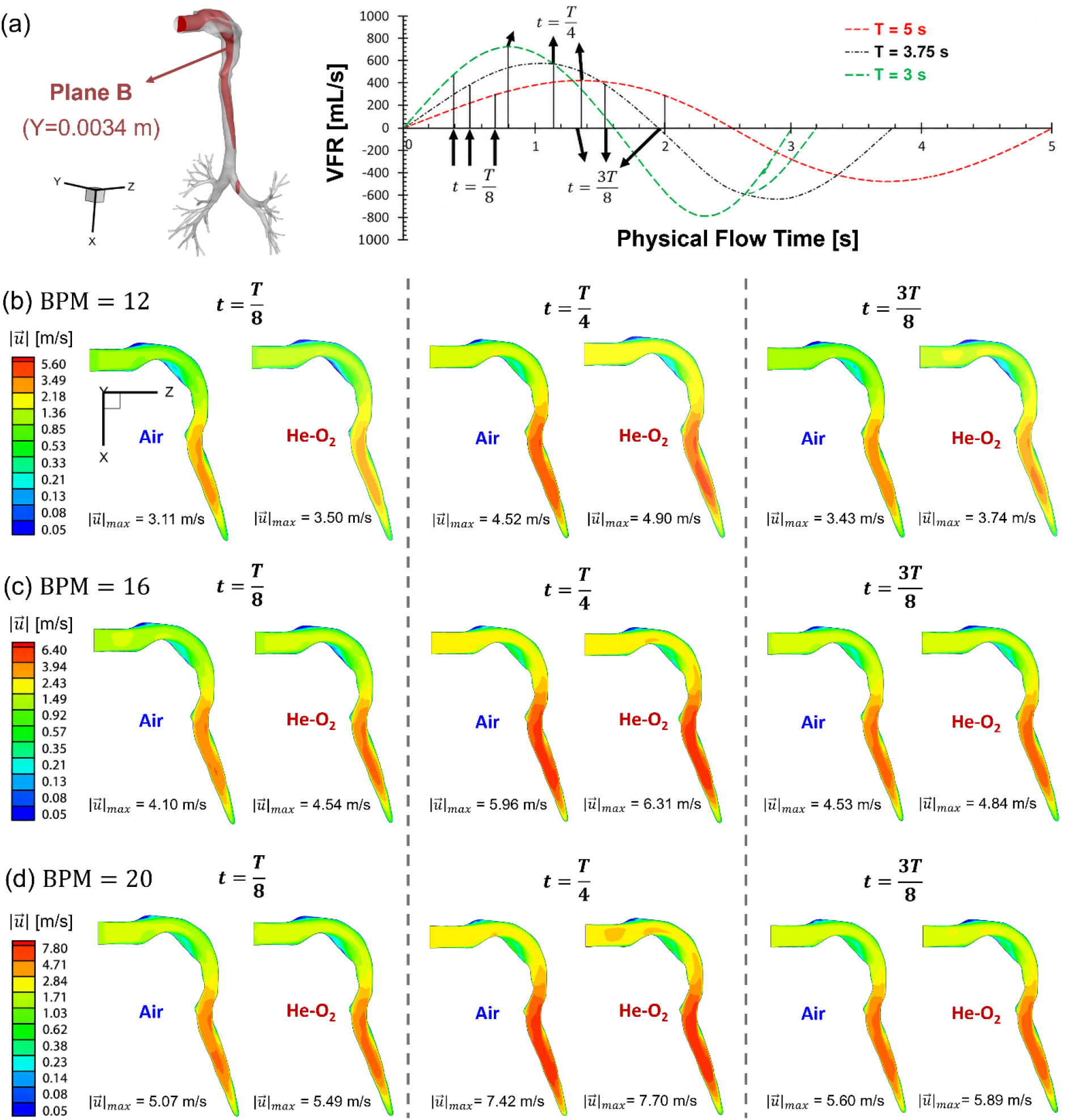
Comparison of velocity magnitude distributions between air and heliox using DLAM on Plane B at the selected inhalation time points 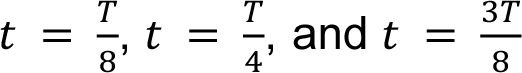: (a) location of Plane B and corresponding breathing waveforms, (b) 12 BPM, (c) 16 BPM, and (d) 20 BPM.

**Fig. 14.**
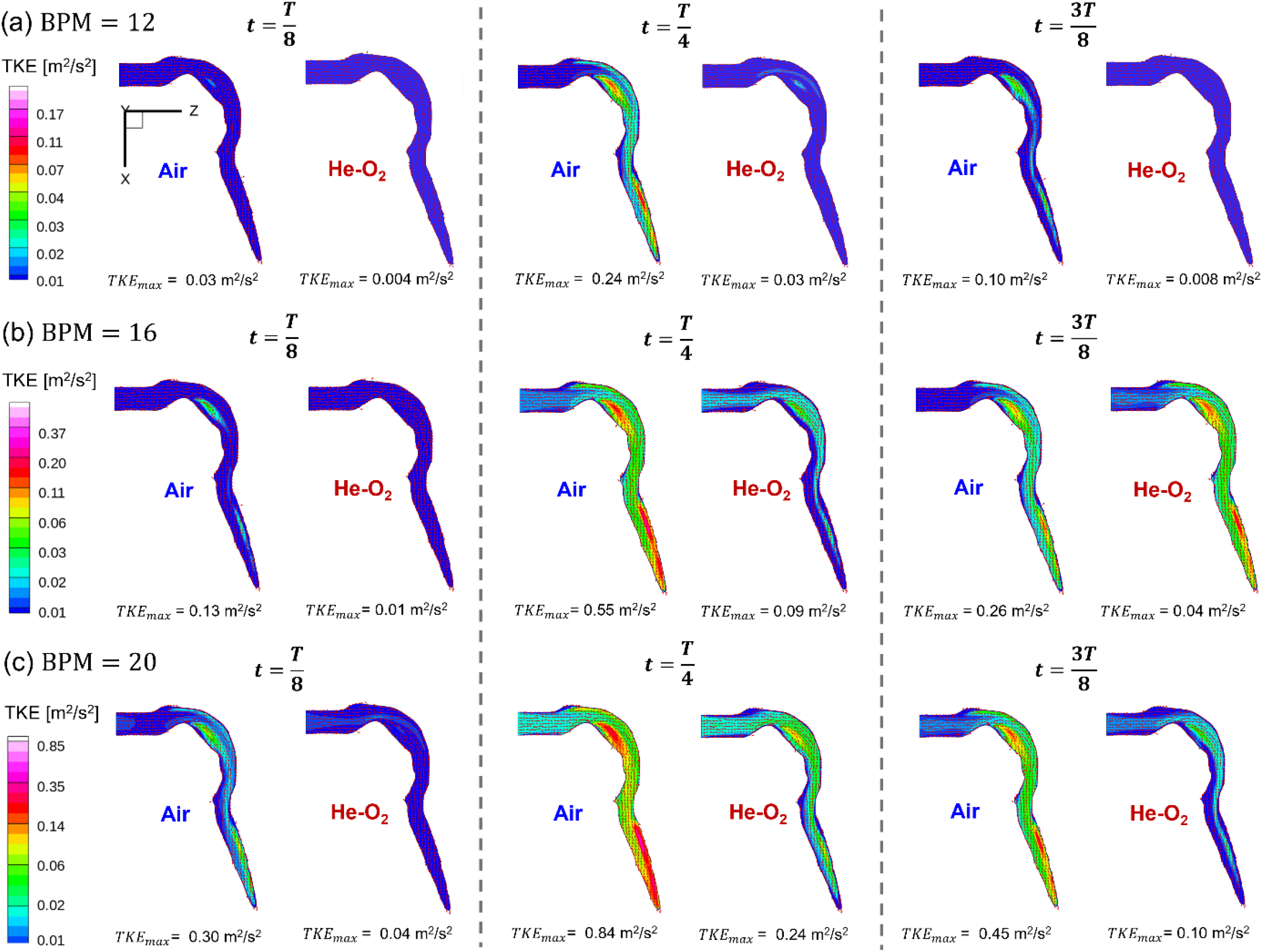
Comparison of turbulence kinetic energy (TKE) distributions and tangential (in-plane) velocity vectors between air and heliox using DLAM on Plane B at 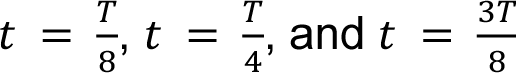: (a) 12 BPM, (b) 16 BPM, and (c) 20 BPM.

Additional comparisons of wall shear stress (WSS) and pressure are shown in **Figs. 15 and 16**, respectively. As shown in **Fig. 15**, the maximum WSS decreases from 7.0 to 6.7 Pa at 12 BPM, from 10.7 to 9.3 Pa at 16 BPM, and from 15.0 to 11.6 Pa at 20 BPM when air is replaced by heliox. Although heliox has a slightly higher dynamic viscosity than air, WSS is determined by both viscosity and the near-wall velocity gradient. The substantially lower density and slightly higher viscosity of heliox reduce the local Reynolds number, which suppresses turbulence and modifies the near-wall momentum transport. The resulting reduction in near-wall velocity gradients outweighs the modest increase in viscosity, leading to lower WSS, particularly at higher breathing frequencies.

**Fig. 15.**
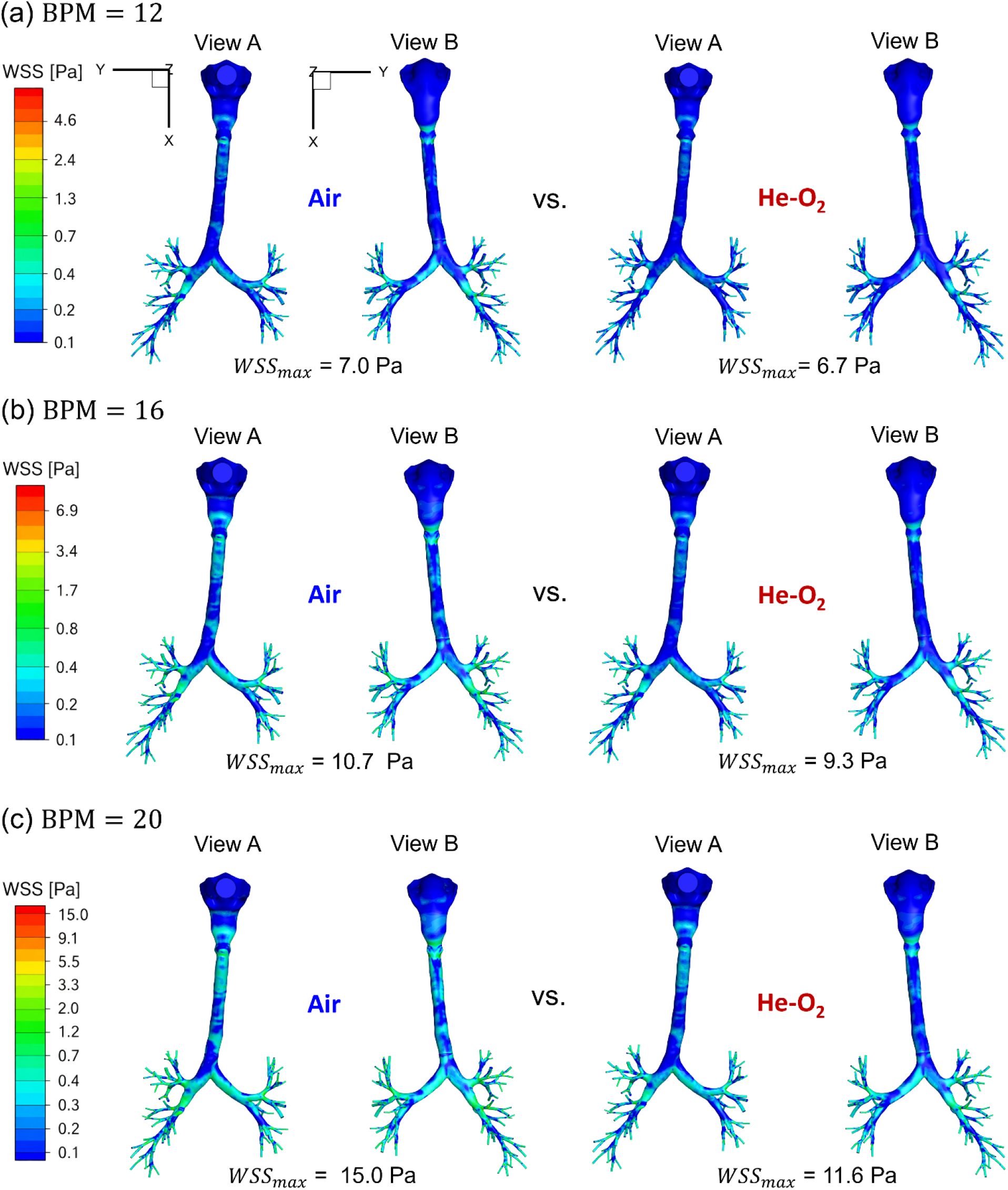
Comparison of wall shear stress (WSS) distributions between air and heliox using DLAM at peak inhalation 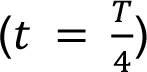: (a) 12 BPM, (b) 16 BPM, and (c) 20 BPM.

**Fig. 16.**
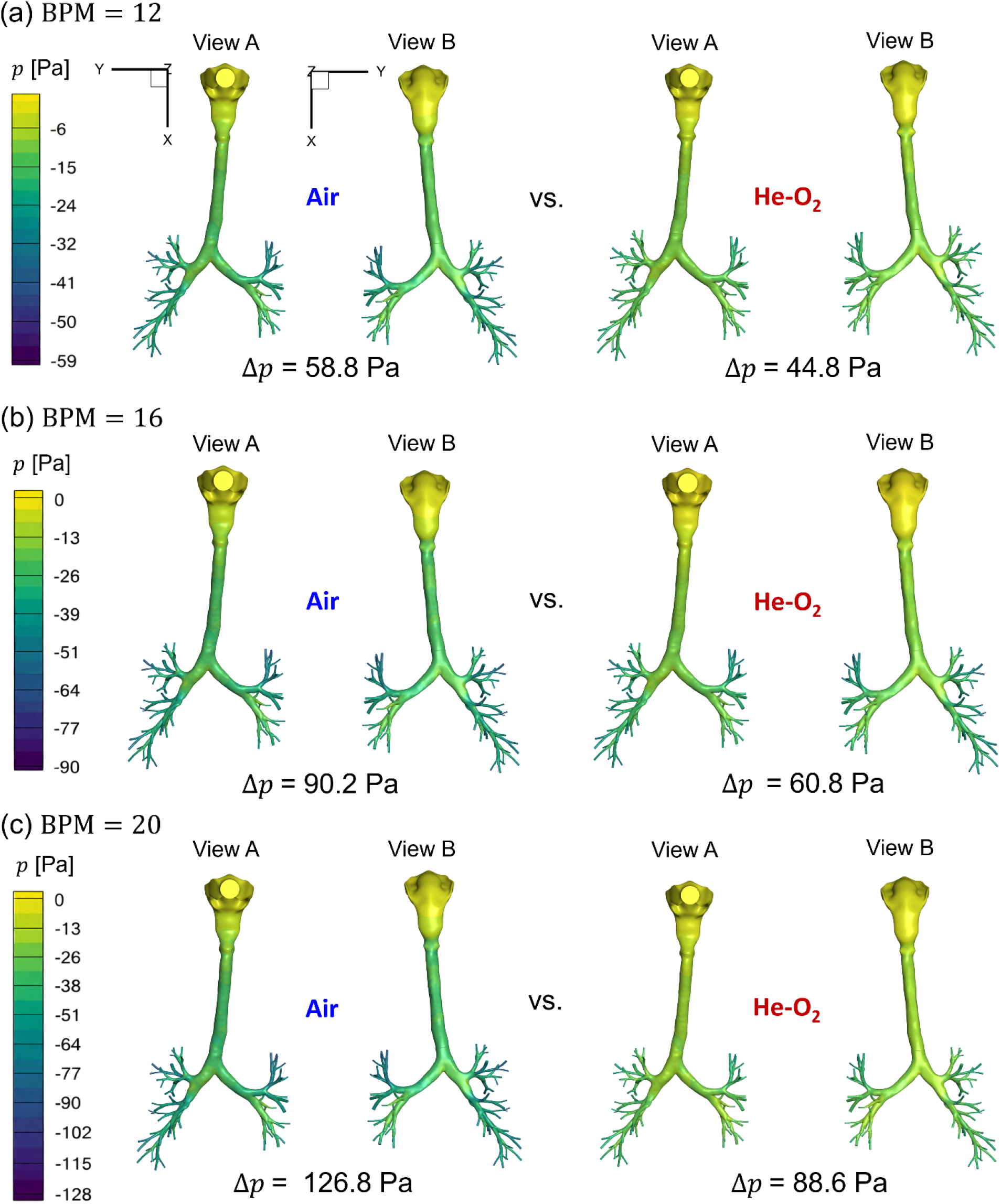
Comparison of gauge pressure distributions between air and heliox using DLAM at peak inhalation 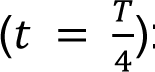: (a) 12 BPM, (b) 16 BPM, and (c) 20 BPM.

Furthermore, a stronger gas carrier effect can be observed for the pressure field in **Fig. 16**. The mouth-to-G9 pressure drop decreases from 58.8 to 44.8 Pa at 12 BPM, from 90.2 to 60.8 Pa at 16 BPM, and from 126.8 to 88.6 Pa at 20 BPM, respectively. Under the same volumetric flow waveform, the much lower density of heliox substantially reduces the inertial pressure requirement and turbulence-related pressure losses, which scale strongly with *ρ*|*u⃗*|^2^. Although its slightly higher viscosity tends to increase the viscous contribution to pressure loss, this contribution is outweighed by the reduction in inertial and turbulence-associated losses. Therefore, the lower WSS and pressure drop predicted with heliox arise from the combined effects of a substantially reduced Reynolds number, weaker turbulence generation, and reduced inertial momentum transport, rather than from a simple reduction in flow velocity.

#### 3.2.2 Influence of breathing frequency on particle transport and deposition

The breathing frequency (i.e., BPM) effect on particle transport and deposition was evaluated at 12, 16, and 20 BPM, while maintaining the same tidal volume of 710 mL (see **Fig. 3** for waveform details). The corresponding breathing-cycle periods were *T* =5, 3.75, and 3 s. Regional deposition fractions and delivery fractions are presented in **Figs. 17-19**, while localized particle deposition patterns can be found in **Figs. 20-22**. The results show that generally increasing BPM results in the same inspired volume to enter over a shorter inhalation interval, resulting in higher instantaneous velocities and greater turbulence theoretically.

**Fig. 17.**
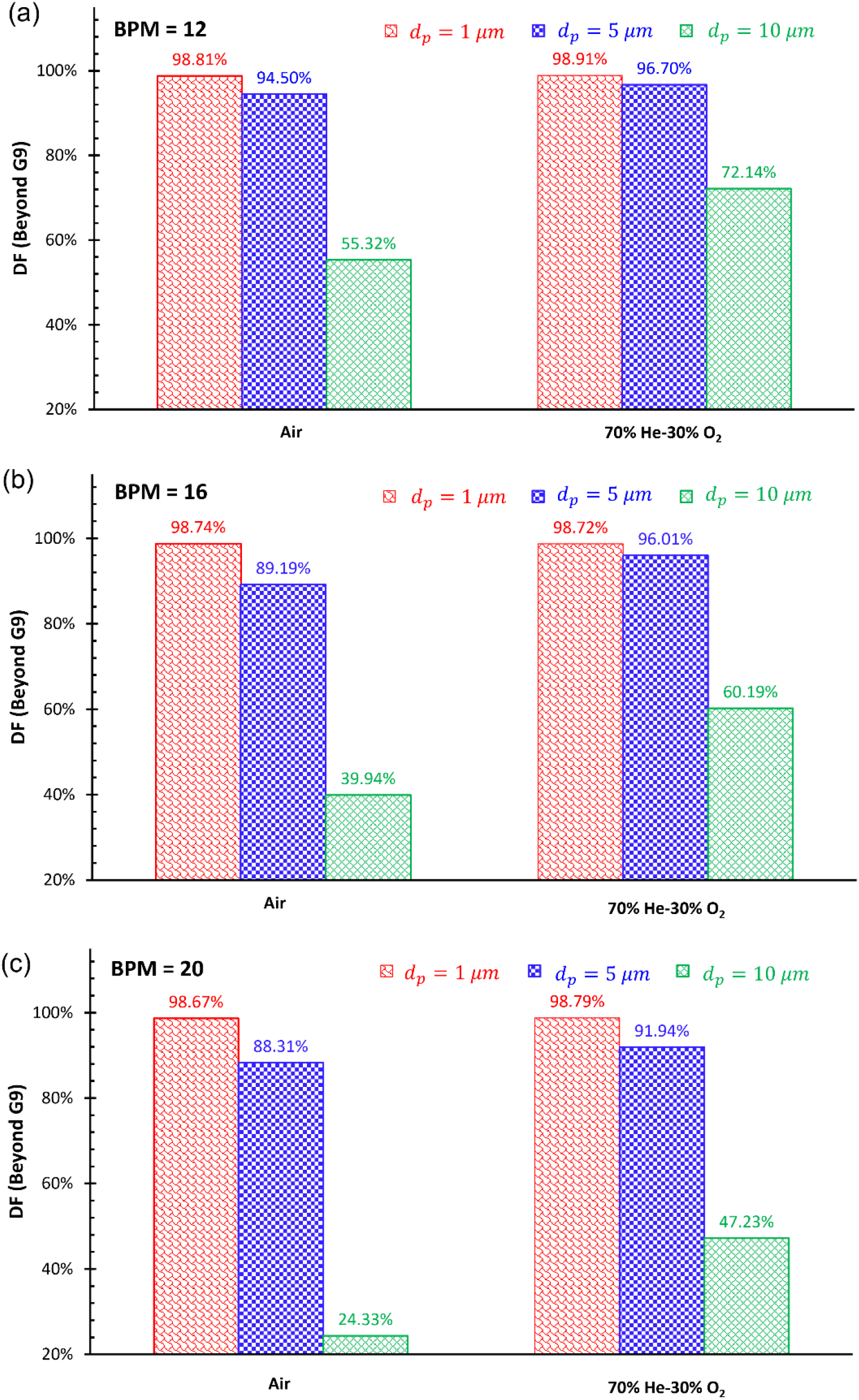
Comparison of beyond-G9 delivery fractions between air and heliox using DLAM for *d_p_* = 1, 5, and 10 *μm* particles at the end of inhalation: (a) 12 BPM, (b) 16 BPM, and (c) 20 BPM.

**Fig. 18.**
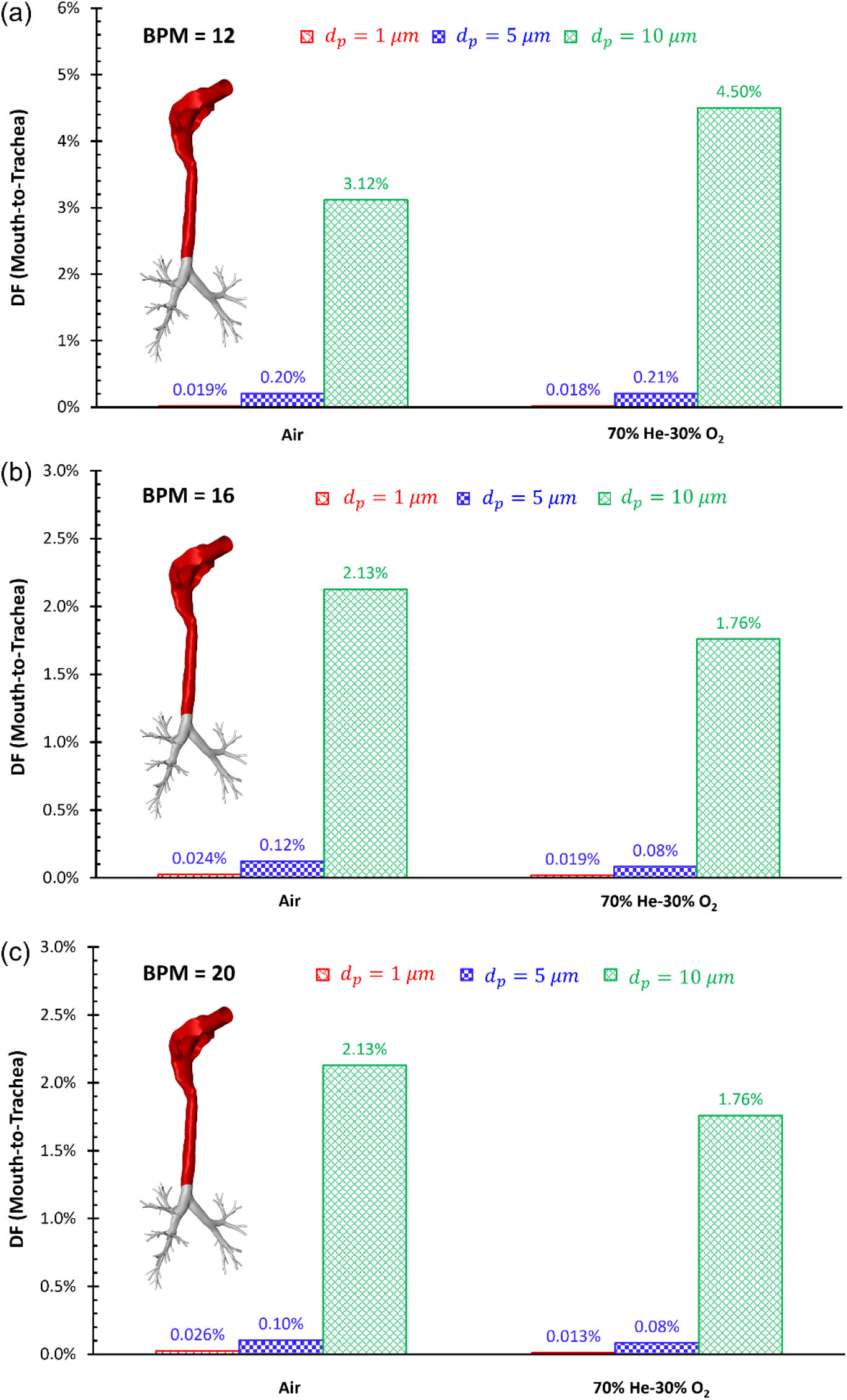
Comparison of mouth-to-trachea regional deposition fractions between air and heliox using DLAM for *d_p_* = 1, 5, and 10 *μm* particles at the end of inhalation: (a) 12 BPM, (b) 16 BPM, and (c) 20 BPM.

**Fig. 19.**
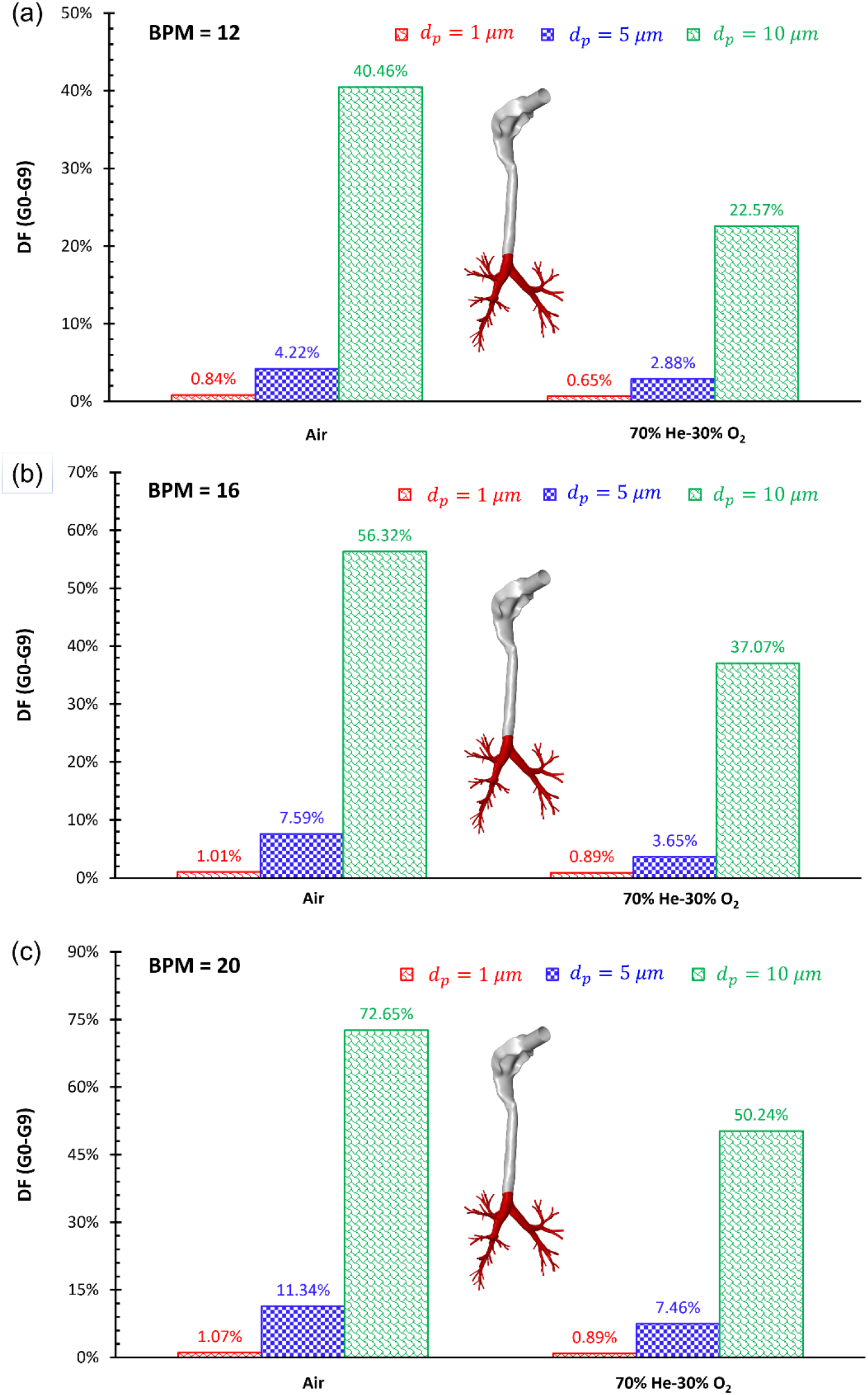
Comparison of G0-G9 regional deposition fractions between air and heliox using DLAM for *d_p_* = 1, 5, and 10 *μm* particles at the end of inhalation: (a) 12 BPM, (b) 16 BPM, and (c) 20 BPM.

**Fig. 20.**
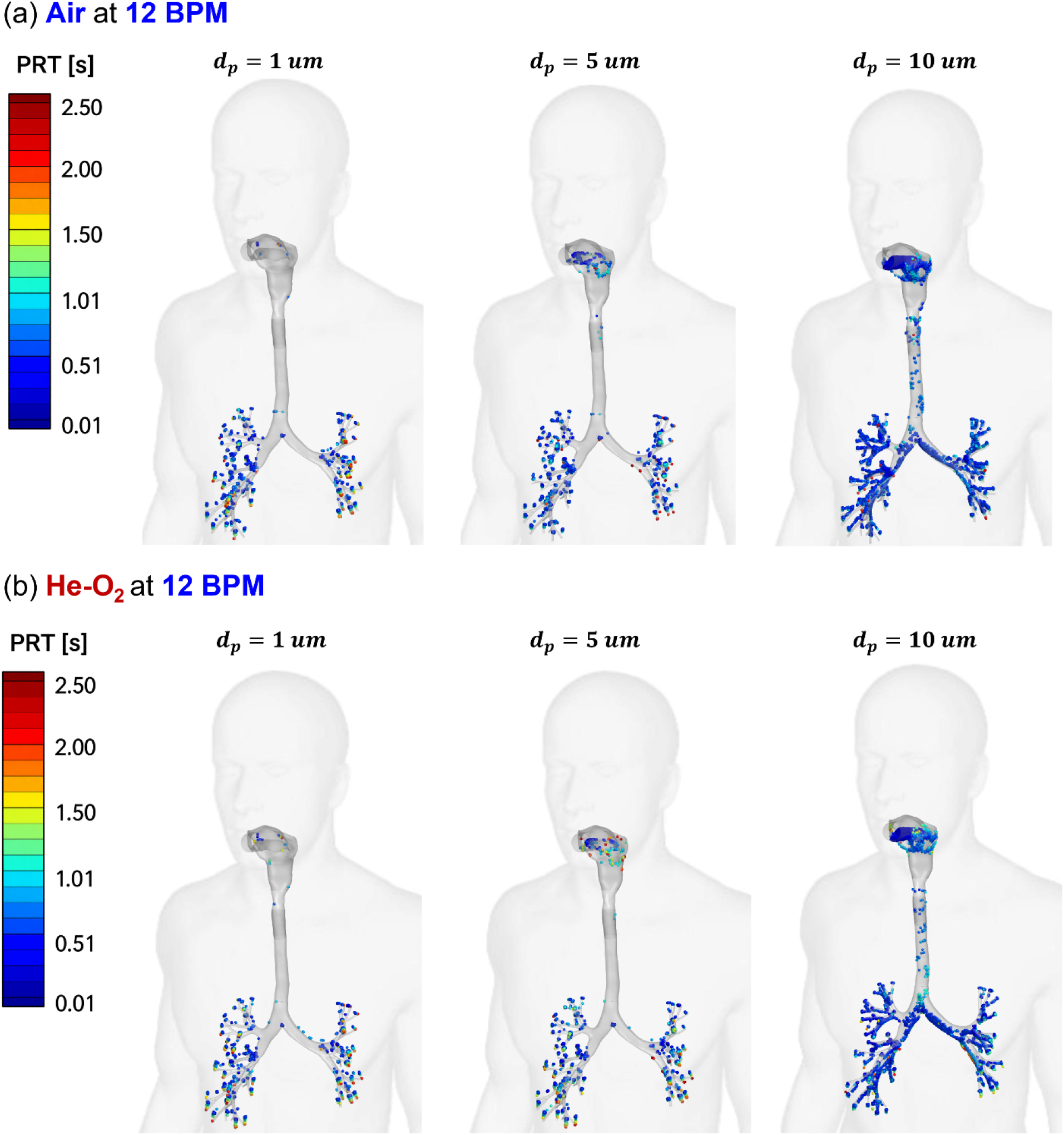
Comparisons of localized particle deposition patterns between air and heliox using DLAM at 12 BPM for *d_p_* = 1, 5, and 10 *μm*: (a) air, and (b) heliox.

**Fig. 21.**
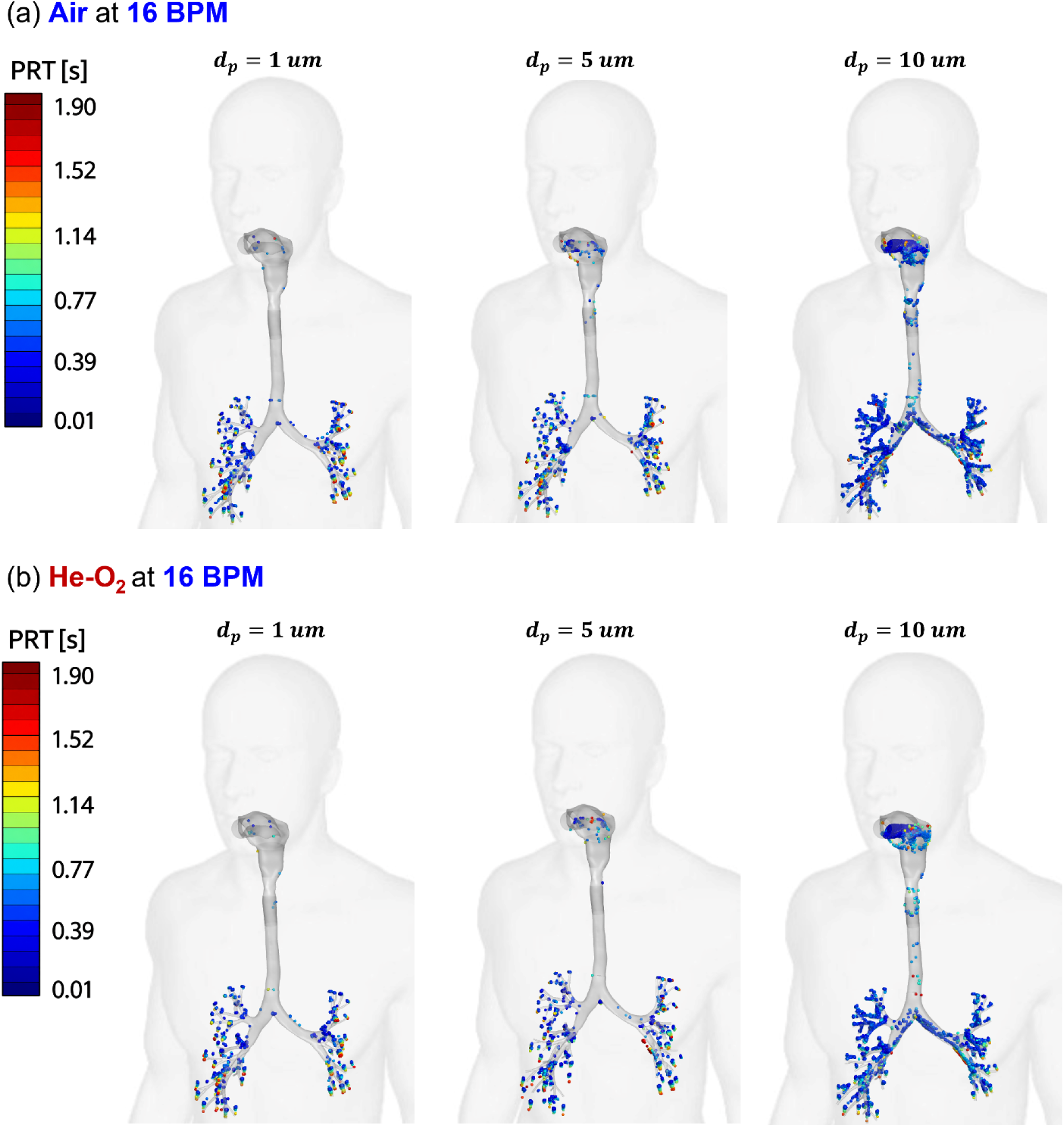
Comparisons of localized particle deposition patterns between air and heliox using DLAM at 16 BPM for *d_p_* = 1, 5, and 10 *μm*: (a) air, and (b) heliox.

**Fig. 22.**
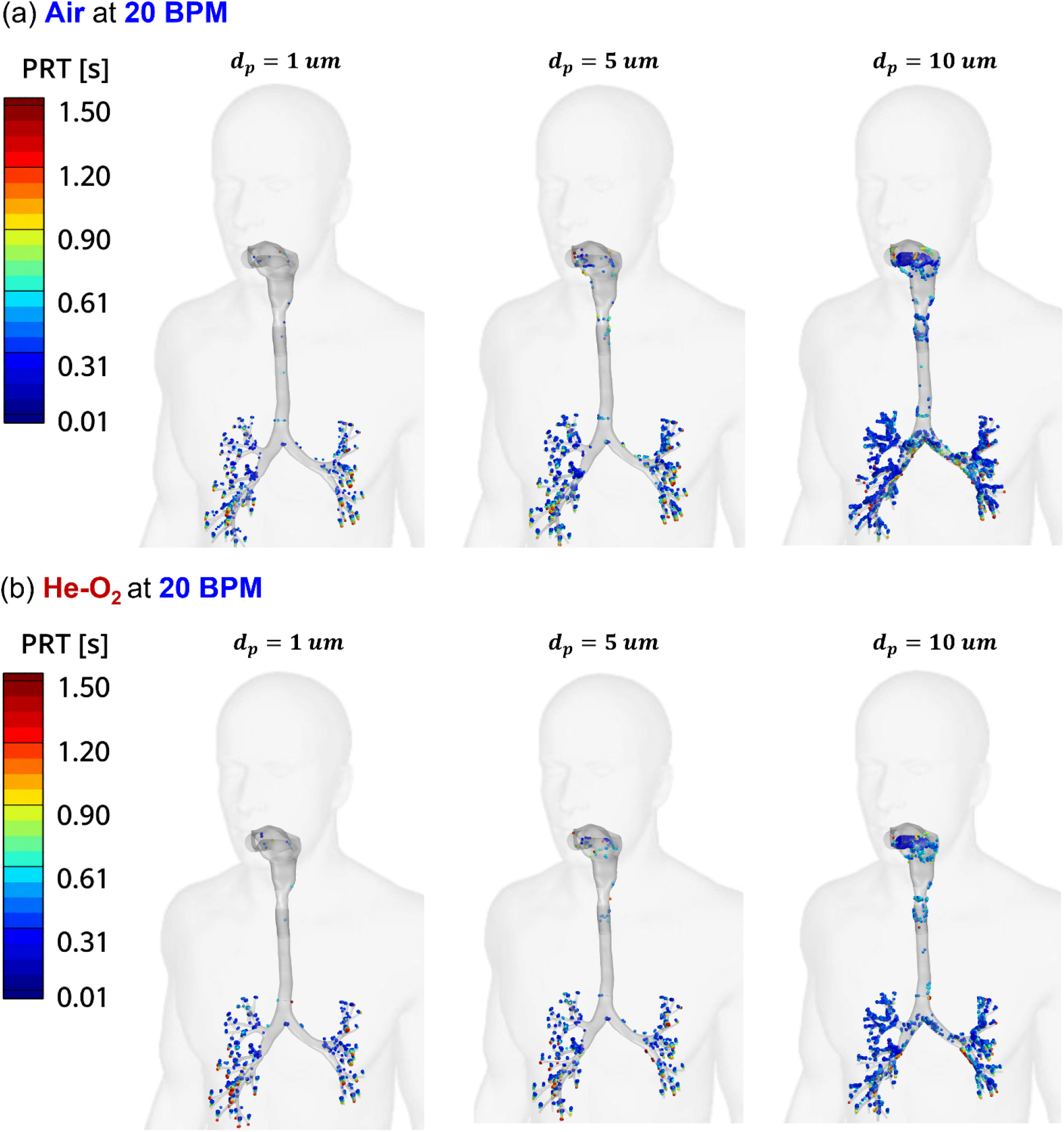
Comparisons of localized particle deposition patterns between air and heliox using DLAM at 20 BPM for *d_p_* = 1, 5, and 10 *μm*: (a) air, and (b) heliox.

For 1 µm particles, the beyond-G9 delivery fraction remains nearly unchanged across the investigated BPMs (see **Figs. 17 (a)-(c)**). The values are 98.81%, 98.74%, and 98.67% for air and 98.91%, 98.72%, and 98.79% for heliox at 12, 16, and 20 BPM, respectively. Since the orders of magnitude of such small variations are comparable to *R*_susp_ reported in **Table 3**, the breathing frequency has no significant influence on beyond-G9 delivery of 1 µm particles under the simulation conditions in this study. In contrast, beyond-G9 delivery decreases significantly with increasing BPM for 5 and 10 µm particles, which are also shown in **Figs. 17 (a)-(c)**. For 5 µm particles, DF decreases from 94.50% to 88.31% with air and from 96.70% to 91.94% with heliox as BPM increases from 12 to 20. For 10 µm particles, it decreases from 55.32% to 24.33% with air and from 72.14% to 47.23% with heliox. Indeed, since the same tidal volume is inhaled over a shorter time duration as BPM increases, the characteristic airflow velocity and local Reynolds number increase. The particle Stokes number also increases, particularly for the 5 and 10 µm particles, making their trajectories less responsive to rapid changes in the carrier-gas streamlines. As a result, particle-flow decoupling becomes stronger in regions of high streamline curvature, airway constriction, and bifurcation, leading to enhanced inertial impaction. At the same time, stronger turbulent fluctuations at higher BPM can enhance cross-stream particle dispersion and increase the probability of collision between particles and airway walls. For larger particles such as the 5 and 10 µm particles, trajectories transported sufficiently close to the airway walls can also undergo interception. Thus, the increased deposition at higher BPM reflects the combined effects of stronger inertial impaction, turbulence dispersion enhanced cross-stream transport, and interception, with inertial impaction expected to become increasingly dominant as particle size increases.

The regional deposition results shown in **Figs. 18 and 19** further identify where these particle losses occur before reaching the G9 terminals. Mouth-to-trachea DFs remain relatively small compared with G0-G9 DFs, whereas G0-G9 DF increases significantly with BPM. For 5 µm particles, G0-G9 DF increases from 4.22% to 11.34% with air and from 2.88% to 7.46% with heliox between 12 and 20 BPM (see **Fig. 19**). For 10 µm particles, the corresponding increases are much larger, from 40.46% to 72.65% with air and from 22.57% to 50.24% with heliox (see **Fig. 19**). The localized deposition patterns in **Figs. 20-22** support this interpretation, demonstrating progressively greater particle accumulation near the lower palate in the oral cavity, pharynx, and bifurcating points, as both BPM and particle size increase. The above-mentioned locations are regions where rapid streamline turning, elevated local velocities, and enhanced turbulent fluctuations collectively increase the chance for particles to strike the airway walls. Therefore, the reduction in beyond-G9 delivery at higher BPM is primarily associated with increased particle losses within the branching G0-G9 airway network, particularly through inertial impaction of the larger particles, with additional contributions from turbulence-enhanced dispersion and interception. Gravitational sedimentation may still contribute to locally low-velocity regions, but the shorter residence time at higher BPM does not favor increased gravitational settling. Accordingly, at a fixed TV, lower breathing frequency promotes beyond-G9 delivery mainly by reducing inertia-dominated particle deposition within the G0-G9 region.

#### 3.2.3 Influence of aerodynamic diameter on particle transport and deposition

Particle size strongly influences the ability of inhaled aerosols to follow the transient carrier gas flow through the human respiratory system. As particle diameter increases, the particle Stokes number increases, indicating stronger particle inertia relative to the characteristic airflow time scale. Larger particles exhibit greater particle-flow decoupling and are more susceptible to inertial impaction at airway bends, constrictions, and bifurcations. Consequently, 1 µm particles respond rapidly to changes in the local gas flow field and remain closely coupled to the carrier-gas streamlines, whereas 5 and especially 10 µm particles exhibit progressively greater particle-flow decoupling. Therefore, larger particles are more susceptible to inertial impaction at locations near the pharynx, glottis, and bifurcating points in TB tree.

This particle size dependence is clearly reflected in the beyond-G9 delivery fractions shown in **Fig. 17**. For air, at 12 BPM, increasing the particle diameter from 1 to 5 and 10 µm decreases the beyond-G9 delivery fraction from 98.81% to 94.50% and 55.32%, respectively. For heliox, the corresponding values decrease from 98.91% to 96.70% and 72.14%. The size effect becomes even more significant at 20 BPM. For air, beyond-G9 delivery decreases from 98.67% for 1 µm particles to 88.31% for 5 µm particles and further to 24.33% for 10 µm particles. For heliox, the corresponding decrease is from 98.79% to 91.94% and 47.23%. Thus, increasing particle size progressively reduces DF beyond G9, and this reduction becomes substantially stronger under the high BPM conditions.

The regional DF results in **Figs. 18 and 19** further demonstrate where the particle-size-dependent deposition increase occurs. The most pronounced increase in deposition is observed within the G0-G9 airways (see **Fig. 19**). At 12 BPM with air, the G0-G9 DF increases from 4.22% for 5 µm particles to 40.46% for 10 µm particles. With heliox, it increases from 2.88% to 22.57%. At 20 BPM, the corresponding increase is from 11.34% to 72.65% for air and from 7.46% to 50.24% for heliox. These results demonstrate that the reduction in beyond-G9 delivery with increasing particle size is primarily associated with enhanced deposition in the TB tree, and a moderate increase in mouth-to-trachea deposition (see **Fig. 18**).

The localized deposition patterns in **Figs. 20-22** provide further spatial distribution evidence of such a particle size dependence. Deposition of 1 µm particles remains sparse throughout the resolved airway for both carrier gases because their low inertia allows them to closely follow the transient gas flow. As particle diameter increases to 5 and 10 µm, deposition becomes progressively concentrated near airway bends, constrictions, and bifurcations, as mentioned in **Section 3.2.2**. It is worth noting that, although such a particle size dependent trend is observed is consistent for both carrier gases, heliox mitigates the loss of larger particles due to its much lower density than air, thereby enhancing the particle delivery into the beyond-G9 region.

#### 3.2.4 Potential physiological effects of heliox as a gas carrier for inhaled medicinal particles

Compared with other potential gas carriers (e.g., H_2_-O_2_ gas mixture) for small airway drug delivery enhancement via inhalation therapy (Yi & Feng, 2026), heliox is biologically inert and does not possess bronchodilating or anti-inflammatory properties. Its effects on respiratory mechanics are governed mainly by its physical properties. The lower density of helium (He) than nitrogen (N_2_) in air leads to reduced turbulence and airway resistance when helium replaces a portion of N_2_ in air, particularly in constricted airways or at high Reynolds number. This mechanism is consistent with the present CFPD results (see **Fig. 16**), where a 70% He-30% O_2_ heliox produced lower pressure drops than air, even under comparable or locally elevated core velocities (see **Section 3.2**). In clinical settings, heliox is primarily utilized to decrease the difficulty of breathing in cases of upper airway obstruction, serving as physical treatment rather than a pharmacological intervention (Hess et al., 2006; Slinger et al., 2019).

For the 70% He-30% O_2_ heliox used in this study, O_2_ concentration inhaled is higher than 21% which is in the ambient air. This O_2_ enrichment may be relevant for patients needing supplemental oxygen. However, the appropriate inspired O_2_ concentration is patient-specific. In patients who are susceptible to oxygen-induced hypercapnia, such as some with severe COPD and chronic CO_2_ retention, oxygen administration should be titrated rather than assumed appropriate based on carrier-gas composition alone (Abdo & Heunks, 2012). Although the current CFPD model with dynamic mesh resolves gas-particle transport, it does not model pulmonary gas exchange, arterial oxygenation, CO_2_ retention, or systemic physiology. Therefore, these results should not be interpreted as demonstrating physiological safety or defining an optimal clinical O_2_ fraction for heliox-based inhalation therapy.

Another practical consideration is that helium can alter the performance of aerosol delivery systems themselves. Bench testing by Li et al. (2024) showed that continuous albuterol delivery with heliox depended strongly on nebulizer type, interface configuration, and heliox composition. In their pediatric model, 80% He + 20% O_2_ and 70% He + 30% O_2_ did not produce identical inhaled doses. Thus, the improved beyond-G9 transport predicted in the present airway model should not be assumed to translate directly into the same delivered drug dose for every inhaler or nebulizer configuration. Device-specific factors, including the nebulizer driving condition, interface fit or leakage, humidification strategy, flow setting, and the selected He-to-O_2_ ratio, should be controlled and experimentally validated before clinical translation.

In summary, although heliox has an extensive history of clinical application, evidence supporting its therapeutic benefit is dependent on specific indications. Slinger et al. (2019) reported that studies of heliox for inducible laryngeal obstruction showed favorable responses. However, the study is insufficient to establish clinical efficacy, emphasizing that physiological plausibility is not equal to therapeutic advantage. Accordingly, the numerical results and findings in this study should be interpreted as mechanistic evidence that heliox can alter airway aerodynamics and inhaled aerosolized medication transport, rather than as proof of clinical superiority.

## 4. Conclusions

This study integrated an experimentally validated Euler-Lagrange CFPD model with a dynamic mesh method (i.e., DLAM), to investigate how airway deformation, carrier gas, breathing frequency, and particle size influence particle transport and deposition from mouth to G9, as well as the delivery into beyond-G9 small airway region. Initial comparisons were made between the static lung airway model (SLAM) and the deformable lung airway model (DLAM) at 12 breaths per minute (BPM). Subsequent analyses using DLAM compared air and heliox (70% He-30% O_2_) at 12, 16, and 20 BPM for particles with 1, 5, and 10 µm in diameter. Key findings are summarized below.

1. Airway deformation kinematics significantly alter the pulmonary gas-particle flow dynamics. Compared to DLAM, SLAM does not account for airway deformation and predict higher localized velocity maxima during inhalation, especially in the glottis region. Additionally, airway deformation redistributes the transient turbulence field instead of uniformly reducing TKE at all locations and time points.
2. The impact of airway deformation kinematics on particle transport and deposition is highly dependent on particle size. The influence on 1 µm particles is negligible. For 5 and 10 µm particles, neglecting airway deformation kinematics (i.e., SLAM) underpredicts delivery beyond G9 and overpredicts deposition within the G0-G9 region, compared with DLAM simulations.
3. DLAM offers a more suitable framework than the conventional SLAM for evaluating gas carrier strategies and inhalation conditions when regional aerosol dosimetry and distal delivery are critical considerations.
4. The carrier gas selection significantly influences pulmonary gas flow dynamics in the DLAM. Compared with air, heliox consistently reduces TKE, WSS, and the pressure drop from the mouth to G9 across all investigated breathing frequencies, while producing comparable or slightly higher localized velocity maxima.
5. The benefit on drug delivery to peripheral lung using heliox as the gas carrier than air increases with particle size. For 1 µm particles, beyond-G9 delivery is nearly equivalent between air and heliox. For 5 and 10 µm particles, using heliox as the gas carrier can reduce mouth-to-G9 airway deposition and enhances beyond-G9 delivery, compared with air.
6. The impact of breathing frequency on particle transport and deposition is also dependent on particle size. At a fixed tidal volume, increasing the breathing frequency has little influence on the beyond-G9 delivery of 1 µm particles but substantially decreases beyond-G9 delivery and increases G0-G9 deposition for 5 and 10 µm particles, with the strongest effect observed for 10 µm particles.
7. Within the investigated conditions, the combination of heliox and a lower breathing frequency with a longer inhalation duration is more favorable for enhancing aerosol delivery beyond G9. Although 1 µm particles exhibit nearly complete beyond-G9 penetration, the present mouth-to-G9 model does not resolve their subsequent deposition in the small airways and alveolar region. Therefore, the optimal particle size for targeted peripheral-lung drug delivery cannot be determined solely from the present results.

## 5. Limitations and Future Work

The present study was designed to isolate the coupled influences of airway deformation, carrier gas, breathing frequency, and particle size while maintaining a computationally tractable model. The limitations of this study and corresponding future work are summarized below:

1. The computational geometry ends at G9. Particles exiting the G9 terminals were assumed to be fully deposited in the unresolved region beyond G9 rather than actual deposition in the small airways or alveolar region. Future studies will extend the airway geometry or couple the current model with validated distal airway and acinar geometries (Zhang et al., 2003).
2. The present simulations were based on one subject-specific airway geometry. Because airway morphology and deformation vary among individuals and disease states, future studies will include multiple healthy and diseased subjects to address the subject variability (Eom et al., 2010; Hasse et al., 2018; Werner et al., 2008; Yin et al., 2013).
3. The parametric study used controlled sinusoidal and quasi-sinusoidal inhalation-exhalation waveforms with a fixed TV to isolate the effects of BPM. Future studies will consider patient- or device-specific inhalation profiles, accounting for variations in inspiratory acceleration, peak flow rate, and breath-hold duration (Dietrich et al., 2026; Yuasa et al., 2015).
4. Particles were assumed monodisperse and spherical with perfect trapping at airway walls. Effects such as hygroscopic growth, evaporation, polydispersity, electrostatic forces, rebound, mucus interaction, and dissolution were not modeled. These factors will be incorporated for formulation-specific prediction (Feng et al., 2021).
5. Only one heliox composition (i.e., 70% He-30% O_2_) was evaluated, and the delivery device was not modeled. Future studies will compare additional He/O_2_ ratios, couple inhaler or nebulizer performance with the 3D airway model, and validate the combined system experimentally (Li et al., 2024). Physiological gas-exchange responses should also be evaluated separately before clinical translation.

## Conflict of Interest

The authors declare no conflict of interest.

## Acknowledgements

This material is based upon work supported by the National Science Foundation under Award Number TI-2234619. The use of Ansys software (Ansys Inc., Canonsburg, PA) as part of the Ansys-CBBL academic partnership is also gratefully acknowledged (Thierry Marchal and Vishal Ganore).

## Declaration of Generative AI Use

Generative AI was used only to assist with language editing and polishing. No AI tools were used to generate scientific results, perform analyses, or develop scientific conclusions. The authors reviewed and approved all AI-assisted edits and will take full responsibility for the accuracy and integrity of the final manuscript.

## Notes

### Competing Interest Statement

The authors have declared no competing interest.

