## Appendix A for "Coupled Effects of Airway Deformation and Carrier Gas on Pulmonary Gas-Particle Flow Dynamics Using an Euler-Lagrange Method with Dynamic Mesh"

### Appendix A: Quantitative Comparison of Pulmonary Flow Velocity Magnitude and Turbulence between SLAM and DLAM for Air and Heliox

Hang Yi and Yu Feng*

School of Chemical Engineering, Oklahoma State University, Stillwater, OK 74078, USA

* **Corresponding Author**

Address: 420 Engineering North, Stillwater, OK 74078, USA

To further quantify how airway deformation kinematics (i.e., DLAM vs. SLAM) influence the flow field and the associated particle transport, area-averaged flow velocity magnitude $|\vec{u}|_{av}$ and area-averaged turbulence kinetic energy $TKE_{av}$ were calculated at selected cross-sections from mouth to generation 2 (G2) (see **Fig. A1**) at three inhalation time stations (i.e., t = T/8, t = T/4, and 3T/8). The inhalation-exhalation waveform with 12 BPM can be found in **Fig. 3**. Specifically, the cross-sections created are listed below, from upstream to downstream in the human respiratory system.

- Plane D is across the region near the mouth opening;
- Plane E is across the larynx;
- Plane F is located above the glottis region;
- Plane C is across the glottis;
- Plane G is below but close to the glottis region;
- Plane H is across the bifurcation site from G0 to G1;
- Plane I is across the bifurcation site from G1 to G2 in the left lobar airways;
- Plane J is across the bifurcation site from G1 to G2 in the right lobar airways.

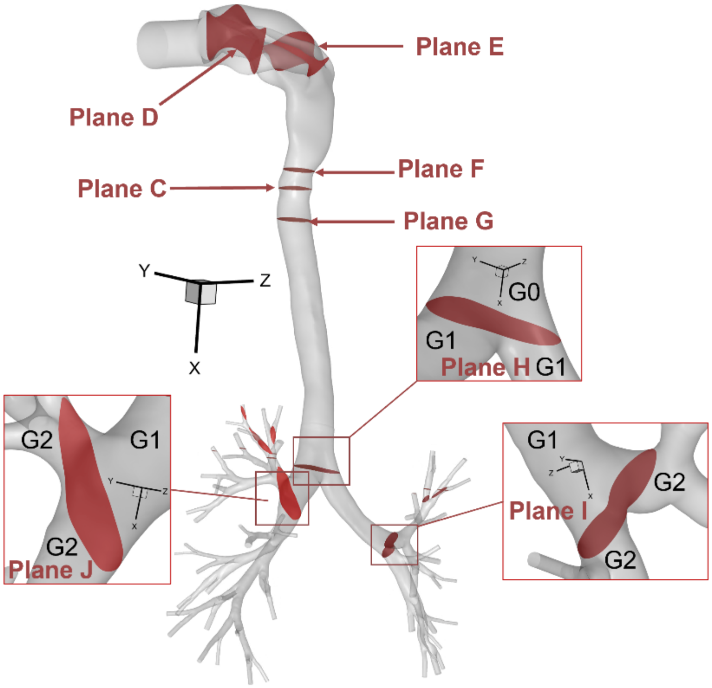


**Figure A1.** Locations of the eight representative cross-sectional planes (i.e., Planes C-J) selected for quantitative comparison of pulmonary gas flow characteristics

The comparisons of $|\vec{u}|_{av}$ and $TKE_{av}$ between SLAM (i.e., without airway deformation) and DLAM (i.e., with airway deformation) with both air and heliox are presented in **Table A1** and **Table A2**, respectively.

**Table A1**: Comparisons of area-averaged $|\vec{u}|_{av}$ [m/s] and $TKE_{av}$ [m²/s²] between SLAM and DLAM on eight representative planes at three different time stations during inhalation (Air).

| | $t = T/8$ | | | | $t = T/4$ | | | | $t = 3T/8$ | | | |
|---|---|---|---|---|---|---|---|---|---|---|---|---|
| | SLAM | | DLAM | | SLAM | | DLAM | | SLAM | | DLAM | |
| | $\|\vec{u}\|_{avg}$ | $TKE_{avg}$ | $\|\vec{u}\|_{avg}$ | $TKE_{avg}$ | $\|\vec{u}\|_{avg}$ | $TKE_{avg}$ | $\|\vec{u}\|_{avg}$ | $TKE_{avg}$ | $\|\vec{u}\|_{avg}$ | $TKE_{avg}$ | $\|\vec{u}\|_{avg}$ | $TKE_{avg}$ |
| Plane C | 2.427 | 2.29E-03 | 1.887 | 1.74E-03 | 3.608 | 1.38E-02 | 2.642 | 1.25E-02 | 2.677 | 7.24E-03 | 2.083 | 5.89E-03 |
| Plane D | 0.577 | 2.01E-03 | 0.552 | 1.75E-03 | 0.881 | 8.43E-03 | 0.842 | 7.38E-03 | 0.654 | 4.03E-03 | 0.624 | 3.53E-03 |
| Plane E | 0.626 | 3.34E-03 | 0.599 | 2.81E-03 | 0.951 | 1.71E-02 | 0.908 | 1.51E-02 | 0.714 | 8.97E-03 | 1.228 | 6.10E-03 |
| Plane F | 2.327 | 2.40E-03 | 2.156 | 1.94E-03 | 3.458 | 1.36E-02 | 3.166 | 1.17E-02 | 2.567 | 7.32E-03 | 2.373 | 6.37E-03 |
| Plane G | 2.258 | 2.35E-03 | 2.150 | 1.90E-03 | 3.361 | 1.55E-02 | 3.203 | 1.75E-02 | 2.490 | 7.60E-03 | 2.370 | 6.74E-03 |
| Plane H | 1.653 | 1.58E-02 | 1.552 | 1.16E-02 | 2.465 | 6.90E-02 | 2.202 | 6.58E-02 | 1.827 | 3.30E-02 | 1.564 | 3.59E-02 |
| Plane I | 1.927 | 1.08E-02 | 1.798 | 6.58E-03 | 2.867 | 5.14E-02 | 2.536 | 4.39E-02 | 2.134 | 2.34E-02 | 1.791 | 2.14E-02 |
| Plane J | 1.563 | 2.70E-02 | 1.468 | 2.37E-02 | 2.311 | 8.48E-02 | 2.051 | 7.89E-02 | 1.719 | 4.69E-02 | 1.453 | 5.04E-02 |

**Table A2**: Comparisons of area-averaged $|\vec{u}|_{av}$ [m/s] and $TKE_{av}$ [m$^2$/s$^2$] between SLAM and DLAM on eight representative planes at three different time stations during inhalation (Heliox).

| | $t = T/8$ | | | | $t = T/4$ | | | | $t = 3T/8$ | | | |
|---|---|---|---|---|---|---|---|---|---|---|---|---|
| | SLAM | | DLAM | | SLAM | | DLAM | | SLAM | | DLAM | |
| | $|\vec{u}|_{avg}$ | $TKE_{avg}$ | $|\vec{u}|_{avg}$ | $TKE_{avg}$ | $|\vec{u}|_{avg}$ | $TKE_{avg}$ | $|\vec{u}|_{avg}$ | $TKE_{avg}$ | $|\vec{u}|_{avg}$ | $TKE_{avg}$ | $|\vec{u}|_{avg}$ | $TKE_{avg}$ |
| Plane C | 2.416 | 2.93E-04 | 1.857 | 2.16E-04 | 3.599 | 2.85E-03 | 2.600 | 2.05E-03 | 2.668 | 1.14E-03 | 2.053 | 8.36E-04 |
| Plane D | 0.561 | 1.07E-03 | 0.536 | 9.59E-04 | 0.850 | 3.31E-03 | 0.812 | 2.95E-03 | 0.630 | 1.68E-03 | 0.601 | 1.50E-03 |
| Plane E | 0.610 | 8.51E-04 | 0.584 | 7.33E-04 | 0.927 | 4.63E-03 | 0.885 | 3.93E-03 | 0.685 | 2.05E-03 | 0.653 | 1.74E-03 |
| Plane F | 2.324 | 3.48E-04 | 2.154 | 2.76E-04 | 3.457 | 3.14E-03 | 3.165 | 2.54E-03 | 2.565 | 1.28E-03 | 2.372 | 1.03E-03 |
| Plane G | 2.249 | 2.55E-04 | 2.144 | 2.05E-04 | 3.347 | 2.64E-03 | 3.188 | 2.17E-03 | 2.482 | 1.03E-03 | 2.366 | 8.31E-04 |
| Plane H | 1.640 | 1.22E-03 | 1.542 | 4.27E-04 | 2.445 | 2.25E-02 | 2.189 | 1.39E-02 | 1.810 | 8.58E-03 | 1.548 | 4.51E-03 |
| Plane I | 1.906 | 1.60E-04 | 1.770 | 6.79E-05 | 2.863 | 6.21E-03 | 2.074 | 4.04E-02 | 2.117 | 1.37E-03 | 1.742 | 4.95E-04 |
| Plane J | 1.549 | 1.15E-03 | 1.449 | 4.40E-04 | 2.313 | 3.65E-02 | 2.505 | 2.35E-03 | 1.717 | 1.21E-02 | 1.471 | 1.55E-02 |
